# A spatial cell atlas of neuroblastoma reveals developmental, epigenetic and spatial axis of tumor heterogeneity

**DOI:** 10.1101/2024.01.07.574538

**Authors:** Anand G. Patel, Orr Ashenberg, Natalie B. Collins, Åsa Segerstolpe, Sizun Jiang, Michal Slyper, Xin Huang, Chiara Caraccio, Hongjian Jin, Heather Sheppard, Ke Xu, Ti-Cheng Chang, Brent A. Orr, Abbas Shirinifard, Richard H. Chapple, Amber Shen, Michael R. Clay, Ruth G. Tatevossian, Colleen Reilly, Jaimin Patel, Marybeth Lupo, Cynthia Cline, Danielle Dionne, Caroline B.M. Porter, Julia Waldman, Yunhao Bai, Bokai Zhu, Irving Barrera, Evan Murray, Sébastien Vigneau, Sara Napolitano, Isaac Wakiro, Jingyi Wu, Grace Grimaldi, Laura Dellostritto, Karla Helvie, Asaf Rotem, Ana Lako, Nicole Cullen, Kathleen L. Pfaff, Åsa Karlström, Judit Jané-Valbuena, Ellen Todres, Aaron Thorner, Paul Geeleher, Scott J. Rodig, Xin Zhou, Elizabeth Stewart, Bruce E. Johnson, Gang Wu, Fei Chen, Jiyang Yu, Yury Goltsev, Garry P. Nolan, Orit Rozenblatt-Rosen, Aviv Regev, Michael A. Dyer

## Abstract

Neuroblastoma is a pediatric cancer arising from the developing sympathoadrenal lineage with complex inter- and intra-tumoral heterogeneity. To chart this complexity, we generated a comprehensive cell atlas of 55 neuroblastoma patient tumors, collected from two pediatric cancer institutions, spanning a range of clinical, genetic, and histologic features. Our atlas combines single-cell/nucleus RNA-seq (sc/scRNA-seq), bulk RNA-seq, whole exome sequencing, DNA methylation profiling, spatial transcriptomics, and two spatial proteomic methods. Sc/snRNA-seq revealed three malignant cell states with features of sympathoadrenal lineage development. All of the neuroblastomas had malignant cells that resembled sympathoblasts and the more differentiated adrenergic cells. A subset of tumors had malignant cells in a mesenchymal cell state with molecular features of Schwann cell precursors. DNA methylation profiles defined four groupings of patients, which differ in the degree of malignant cell heterogeneity and clinical outcomes. Using spatial proteomics, we found that neuroblastomas are spatially compartmentalized, with malignant tumor cells sequestered away from immune cells. Finally, we identify spatially restricted signaling patterns in immune cells from spatial transcriptomics. To facilitate the visualization and analysis of our atlas as a resource for further research in neuroblastoma, single cell, and spatial-omics, all data are shared through the Human Tumor Atlas Network Data Commons at www.humantumoratlas.org.

## INTRODUCTION

Pediatric solid tumors can arise from mesodermal, endodermal and ectodermal lineages during development and retain the molecular and cellular features of their embryonic origins^1^. Single-cell and single-nucleus RNA-seq (sc/sn-RNA-seq) of pediatric solid tumors have showed that they contain heterogeneous populations of malignant cells representing different developmental stages within particular embryonic lineages^2–11^. For example, embryonal rhabdomyosarcomas are composed of malignant cells resembling multiple stages of skeletal muscle differentiation^9–11^. Beyond this developmental hierarchy, there are intra- and inter-tumor variability in the composition of the non-malignant tumor microenvironment (TME). Signaling between the malignant and non-malignant cells in tumors can impact treatment response, including to chemoimmunotherapy.

Neuroblastoma is a pediatric solid tumor of the developing sympathoadrenal lineage. The sympathoadrenal lineage is specified with the formation of proliferating sympathoblasts during the first wave of neural crest migration (E8.5-E9.5 in mice and PCD 27-30 in humans)^6,12,13^. During the early stages of sympathoadrenal development, sympathoblasts produce post-ganglionic sympathetic neurons that are either adrenergic or cholinergic^14^. Later, these sympathoblasts produce Schwann cell precursors (SCPs), a multipotent cell population that can differentiate into Schwann cells, multiple mesenchymal cell types, and adrenergic chromaffin cells of the adrenal medulla^15–18^. Both intrinsic and extrinsic factors regulate the specification and differentiation of adrenergic and mesenchymal cells from sympathoblasts^14^. Findings in neuroblastoma cell lines mirror the presence of both adrenergic and mesenchymal cell populations in the developing adrenal gland, with the presence of two interconvertible epigenetic states in malignant cells, termed the adrenergic (ADRN) and mesenchymal (MES) cells^19–22^. In addition to the striking heterogeneity within individual tumors, neuroblastomas also exhibit patient-to-patient variability with broad clinical outcomes. For example, *MYCN* amplification and age at diagnosis are the two of the most significant predictors of outcome, with survival rates 5 to 10 times higher in infants than in adolescents or young adults^23–25^. While patients with metastatic disease often have a poor outcome despite intensive multi-modal therapy^23,26^, a rare subset of neonates and infants with widely metastatic neuroblastomas experience spontaneous disease regression and have excellent outcomes^27,28^. Moreover, tumors with well-differentiated adrenergic malignant cells or those dominated by a non-malignant Schwannian stroma are associated with more favorable patient outcomes; these findings have led to the incorporation of histologic status in clinical risk stratification^29^. Finally, clinical success using antibodies and chimeric antigen receptor (CAR) T cell therapy directed towards GD2 ganglioside demonstrates a potential role for engaging the immune system for effective treatment of neuroblastoma^26,30–34^. Taken together, these findings indicate that there is a complex interplay between intra-tumor cellular heterogeneity in both the malignant and non-malignant compartment, inter-patient variability, and outcome in neuroblastoma.

Multiple scRNA-seq studies have provided important insights into the cellular heterogeneity of neuroblastomas and the normal developing sympathoadrenal lineage^2,4,6,8,12,35,36^. However, these prior studies have focused on smaller cohorts of 8-20, and, thus far, have led to contradicting conclusions regarding the presence or absence of malignant neuroblastoma cells resembling cells of the developing sympathoadrenal lineage. Moreover, the spatial distribution of malignant cell states, and their relation to the TME, remain to be more comprehensively analyzed in neuroblastoma.

Here, we report a comprehensive cellular atlas of neuroblastoma from the Human Tumor Atlas Pilot Project (HTAPP)^37^, which spans 55 tumors from 51 pediatric patients, and encompasses the full clinical spectrum of neuroblastoma. We identified three major populations of malignant cells with features of proliferating sympathoblasts (SYMP), adrenergic neurons (ADRN), and mesenchymal cells (MES). These cell populations were not uniformly distributed across our cohort, and we introduce methylation profiling, which identified four groups of neuroblastoma that differed by the degree of malignant heterogeneity. Moreover, two methylation groups had significantly worse disease outcomes, suggesting that methylation profiling may have utility as a potential risk stratifier of disease. Non-malignant cells in the neuroblastoma TME also exhibited intra- and inter-tumor heterogeneity, and spatial analysis demonstrated a compartmentalized structure where immune cells were more likely to be adjacent to MES tumor cells and separate from SYMP and ADRN tumor cells. Moreover, we identified spatially distinct expression programs in immune cells that depended on their relative adjacency to malignant cells. Our study sheds new light on malignant expression programs in neuroblastoma and an atlas for the future studies its intra- and inter-tumoral heterogeneity.

## RESULTS

### A cellular, genomic and spatial atlas of neuroblastoma

We built a tumor atlas for neuroblastoma as part of HTAPP, a pilot for the human tumor atlas network (HTAN)^37^, by collecting samples from two pediatric cancer centers (St. Jude Children’s Research Hospital and Dana-Farber Cancer Institute). In total, we selected 55 tumors from 51 patients that met the criteria for sc/snRNA-seq and spatial analyses. These neuroblastomas span the clinical breadth of disease, including age, international neuroblastoma staging scale (INSS) stage, and histology (neuroblastoma and ganglioneuroblastoma) (**Figure 1A** and **Table S1**). They included neuroblastoma samples with *MYCN* amplification (n=12) and *ALK* gain-of-function mutations (n=9; **Figure 1A** and **Table S2**). Twenty-seven samples (predominantly stage 3 or 4) were obtained in the midst of treatment with chemotherapy, nine of which received chemo-immunotherapy with an anti-GD2 antibody^38,39^. For patient HTAPP-312, we analyzed samples before and after chemotherapy treatment; HTAPP-800 had two different sites, and HTAPP-811 had three timepoints during chemo-immunotherapy treatment.

**Figure 1.**
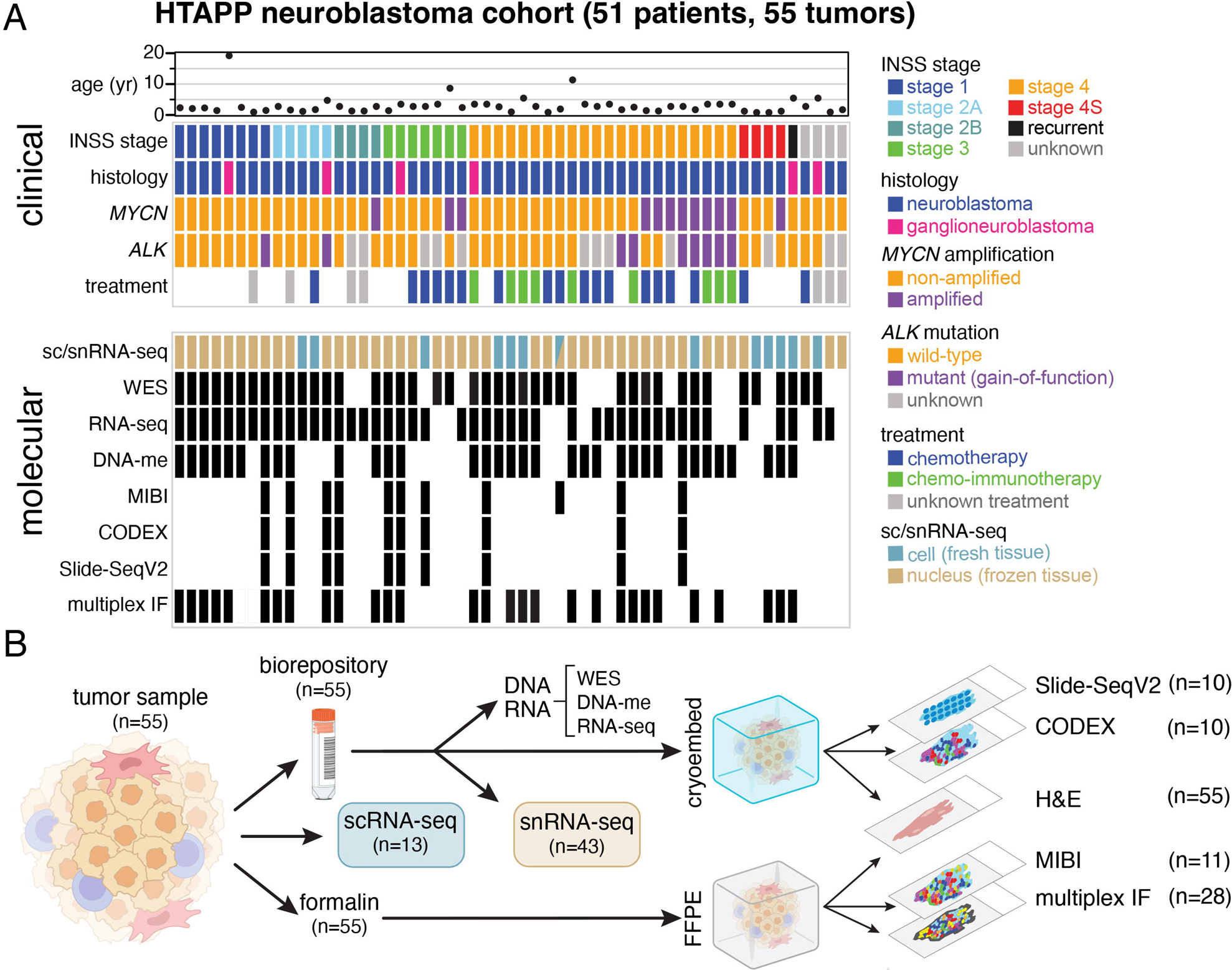
HTAPP neuroblastoma study design and sample cohort. (A) Clinical, histologic, and molecular features of the HTAPP neuroblastoma dataset (n=55). Sequencing and spatial technologies applied are indicated. (B) Sample workflow. Fresh, frozen, and fixed tissue from neuroblastoma tumors were from two institutions, St. Jude Children’s Research Hospital and Dana-Farber Cancer Institute. Fresh or frozen tissue were dissociated for single-cell or single-nucleus RNA-sequencing, respectively. Additionally, a subset of tumors underwent deep spatial profiling using spatial proteomic (MIBI and CODEX) or spatial transcriptomic (Slide-SeqV2) methods. Abbreviations: INSS, international neuroblastoma staging system; WES, whole exome sequencing; DNA-me, DNA methylation; MIBI, multiplexed ion beam imaging; CODEX, co-detection by indexing; FFPE, formalin-fixed paraffin-embedded; IF, immunofluorescence.

To elucidate the cellular heterogeneity across our cohort, we profiled 13 fresh tumor specimens by scRNA-seq, 43 frozen specimens by snRNA-seq, and one sample (HTAPP-656-SMP-7481) by both (**Figure 1A** and **Table S1, Supplemental Information**). SnRNA-seq applied to frozen samples allowed us to exploit previously harvested and archived samples from both centers, thus increasing the number of patient samples available for study from a rare pediatric solid tumor like neuroblastoma and include tumors for which clinical outcomes are available. We successfully generated data from samples that were banked up to ten years prior to snRNA-seq. We also performed whole exome sequencing (WES) of tumor and germline DNA (n=43), bulk RNA-seq (n=44), and DNA methylation (Illumina 850k BeadChip arrays; n=34) (**Figure 1A**, **Tables S1**, **S2, Supplemental Information**).

For the spatial atlas, tumors were apportioned in parallel for formalin fixation and paraffin embedding (FFPE) and for cryoembedding and cryosectioning (**Figure 1B, Supplemental Information**). From the FFPE specimens, all 55 samples had histologic staining with hematoxylin and eosin (H&E; initially used for clinical diagnostic purposes), 28 of the tumors were further analyzed by multiplexed immunofluorescence, and 11 were analyzed by multiplexed ion beam imaging (MIBI)^40,41^ (**Figure 1B, Supplemental Information**). Serial sections from 10 of the cryoembedded tumors were profiled by the spatial transcriptomics method Slide-seqV2^42,43^ and by the spatial proteomics method CO-Detection by indexing (CODEX)^44^ (**Figure 1B, Supplemental Information**). We selected the 10-11 tumors for spatial profiling to span most INSS stages and by an anatomic pathologist’s review to avoid samples with large regions of necrosis or calcification. Notably, 10 of 11 patient tumors were used across all three spatial platforms (Slide-SeqV2, CODEX and MIBI), to compare and cross-validate the different assays, and facilitate future development of computational methods (**Figure 1A,B**, **Table S1**). All raw, processed, and annotated data from this study are available on the HTAN data commons (https://humantumoratlas.org/explore).

### Neuroblastomas have three major malignant cell states

The atlas included 530,055 high-quality^45^ cell and nucleus profiles (84,769 cells; 445,286 nuclei; **Table S3, Supplemental Information**). These profiles were processed uniformly^45,46^ (**Figure S1A**; **Supplemental Information**), combined using integrative non-negative matrix factorization (iNMF), and co-embedded into a shared low-dimensional latent space^47^ (**Figure 2A** and **S1B,C**). Consistent with our previous studies^45,48^, snRNA-seq profiles had significantly fewer expressed genes (1,373 genes/nucleus) than scRNA-seq (2,101 genes/cell) (p < 2.2×10^−16^), lower proportion of mitochondrial reads (0.3% versus 4.4%; p < 2.2×10^−16^) (**Figure S1D-F**), and lower scores for two signatures of dissociation-induced stress^49,50^ (**Figure S1G,H**; p < 2.2×10^−16^). The iNMF integrated profiles partitioned into 20 clusters (**Figure 2A-C**; **Table S4**), annotated by marker expression: five immune cell clusters (*e.g.*, CD45 (*PTPRC*), **Figure 2C** and **S2A**), three endothelial cell clusters (*e.g.*, CD31 (*PECAM1*), **Figure 2C** and **S2B**), and 12 clusters that express genes of the developing sympathoadrenal lineage (*e.g.*, NCAM (*NCAM1*), **Figure 2B,C** and **S2C**). The 12 sympathoadrenal clusters were further partitioned into three groups: seven clusters expressing genes representing different stages of sympathetic neuron differentiation (*NRG3, NTRK3, DBH, ERBB4, CHGB*) (**Figure 2C**), consistent with the adrenergic (ADRN) gene expression signature found in N-type neuroblastoma cell lines^14,51,52^; four clusters expressing genes found in mesenchymal cells derived from the sympathoadrenal lineage (*FN1, ACTA2, COL1A1, EGFR*) (**Figure 2C**), consistent with the mesenchymal (MES) gene expression signature found in S-type neuroblastoma cell lines^14,51–53;^ and one cluster resembling proliferating sympathoblasts (*MKI67, TOP2A*) (**Figure 2C**), which lie at the junction between the sympathetic neurons and the mesenchymal cells consistent with their developmental competence to produce both cell lineages (**Figure 2B**).

**Figure 2.**
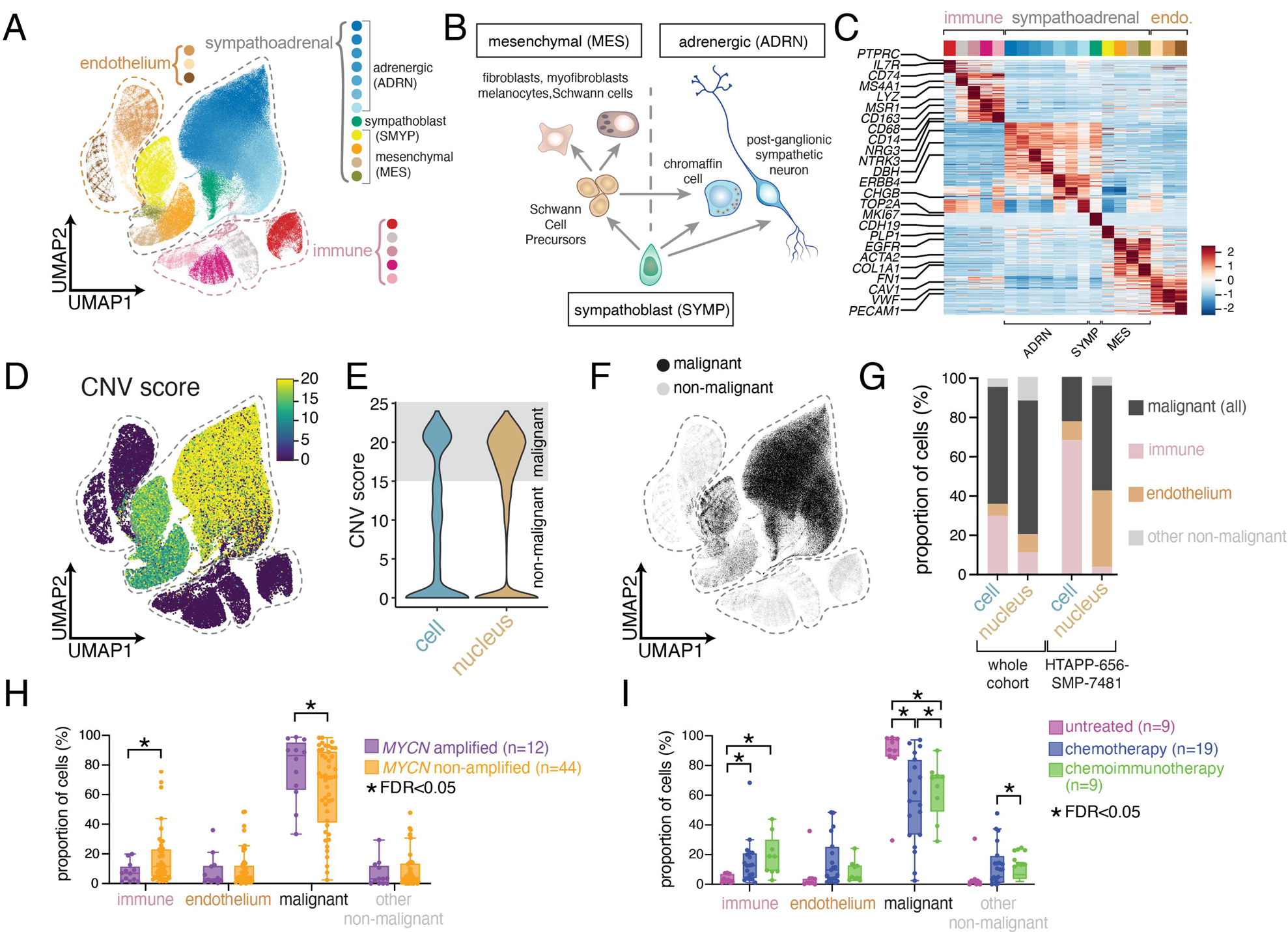
Single-cell and nucleus RNA-sequencing of 55 neuroblastoma tumors. (A) UMAP plot showing integrated data from both scRNA-seq (n=13 tumors; 84,769 cells) and snRNA-seq (n=43 tumors; 445,286 nuclei). Twenty clusters were identified, which were annotated as endothelial (n=3 clusters), immune (n=5 clusters), or sympathoadrenal (n=12 clusters). (B) Schematic of sympathoadrenal lineages within the developing adrenal gland. Sympathoblasts are a proliferative sympathoadrenal progenitor, which differentiate to generate either: (i.) cells of the mesenchymal (MES) lineage, which are derived from multipotent Schwann cell precursors (SCPs), or (ii.) cells of the adrenergic (ADRN) lineage, which consist of postmitotic adrenergic cells. (C) Heatmap showing relative expression of the top 30 genes for each joint cluster. Expression is colored based on scaled normalized value (z-score). (D) UMAP plot showing copy number variant (CNV) scores following inference of copy-number alteration. (E) Violin plot comparing the CNV score from scRNA-seq and snRNA-seq data. (F) UMAP plot as in (A) colored based on the presence or absence of inferred copy number alterations. (G) Cell type composition after coarse annotation of the transcriptomic atlas, comparing all single-cell RNA-seq data and all single-nucleus RNA-seq data, or comparing one sample (HTAPP-656-SMP-7481) that was processed by both single-cell and single-nucleus RNA-sequencing. (H-I) Relative proportion of immune, endothelium, malignant, and non-malignant cells/nuclei for each tumor comparing MYCN status (H) or treatment status (I). The comparison of treatment impact in heterogeneity (HI was restricted to only intermediate-risk or high-risk samples. Bars with a * demarcate credibly significant differences, as calculated by Bayesian composition analysis^88^ using a false discovery rate (FDR) of 0.05. Abbreviations: UMAP, uniform manifold approximation and projection; ADR, adrenergic; S, sympathoblast; MES, mesenchymal; SCP, Schwann cell precursor; CNV, copy number variation; FDR, false discovery rate.

We distinguished malignant from non-malignant profiles by their inferred copy number alterations (CNAs) from sc/snRNA-seq profiles^45,54^, identifying 344,124 malignant cell profiles (64.9% of total), all restricted to the sympathoadrenal clusters (**Figure 2D-F** and **S2D,E**). Each malignant cell subset (SYMP, ADRN, MES) had similar patterns of inferred CNAs (**Figure S2E**). The proportion of immune cells was higher in scRNA-seq, while snRNA-seq captured a higher fraction of malignant and non-malignant cells with sympathoadrenal lineage signatures, consistent with previous reports in other tumors and healthy tissues^45,48,55^ (**Figure 2G**). This was also reflected in the one sample profiled by both scRNA-seq and snRNA-seq, HTAPP-656-SMP-3481^45^ (**Figure 2G** and **S2F**).

We validated the proportion of SYMP, ADRN and MES cells in neuroblastoma by immunohistochemical staining for MKI67 (sympathoblasts), PHOX2B (adrenergic) and VIM (mesenchymal) for 15 tumors (**Figure S2G**, **Table S5**). We found good concordance between the IHC and snRNA-seq (**Figure S2H,I;** r^2^=0.77-0.96), consistent with the agreement observed in other tumor types^55^. We have developed an online interactive visualizer to help interrogate the single cell/nucleus atlas, along with individual sample CNA inference heatmaps (https://viz.stjude.cloud/community/human-tumor-atlas-network-consortium~6).

### Tumor composition shifts associated with MYCN amplification or therapy

Next, we used a Bayesian composition model to compare the proportions of cells (malignant, immune, endothelial, other non-malignant) with or without *MYCN* amplification or treatment (**Figure 2H,I, Supplemental Information**). *MYCN*-amplified neuroblastomas had a reduction in the proportion of immune cells within tumors (FDR<0.05), consistent with prior reports^56–58^. Conversely, intermediate and high-risk tumors treated with either chemotherapy or chemoimmunotherapy had a statistically credible reduction in malignant cell proportion and an increase in the proportion of immune cells (FDR<0.05) (**Figure 2I**).

### Adrenergic and mesenchymal cell heterogeneity in neuroblastoma malignant cells

To further characterize the heterogeneity within malignant cell/nuclei profiles, we reintegrated only the 50,532 malignant cell and 293,592 malignant nucleus profiles and partitioned them into 12 clusters (**Figure 3A-C**; **Table S5**). We identified three clusters enriched for mesenchymal (MES) genes, seven clusters enriched for adrenergic genes (ADRN) and one proliferating sympathoblast (SYMP) cluster (**Figure 3A-C**).

**Figure 3.**
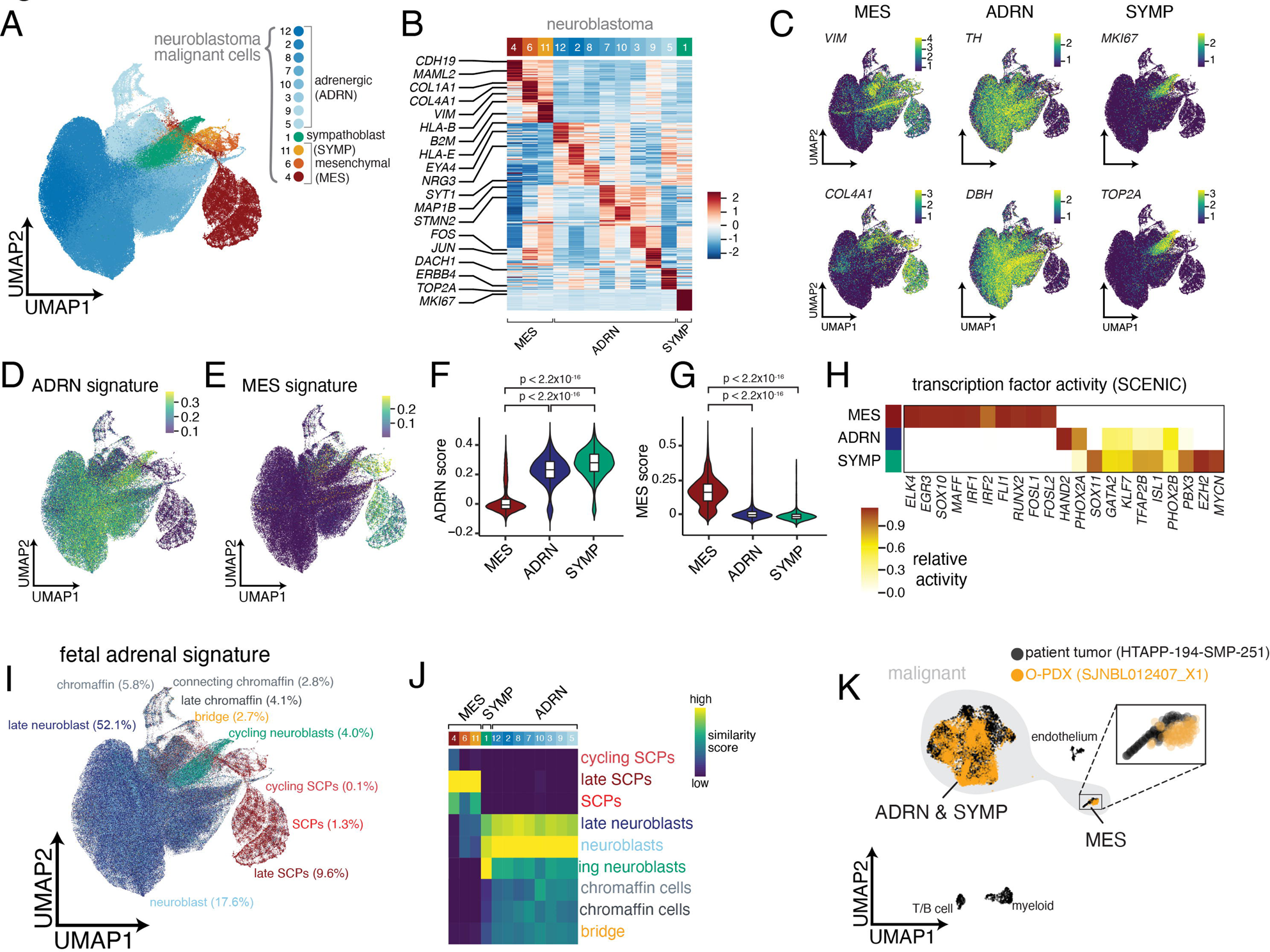
Analysis of malignant cell-specific programs of neuroblastoma. (A) UMAP plot showing integrated data of malignant cells/nuclei from scRNA-seq (n=13 tumors; 50,532 cells) and snRNA-seq (n=43 tumors; 293,592 nuclei) clustered by joint cluster. Twelve clusters were identified which were annotated as adrenergic (n=8 clusters), mesenchymal (n=3 clusters), or sympathoblast (n=1 cluster). (B) Heatmap showing relative expression of the top 30 genes for each malignant cluster. Expression is colored based on scaled normalized value (z-score). (C) UMAP plots of malignant cells/nuclei, colored based on expression of mesenchymal markers (*VIM*, *COL4A1*), adrenergic markers (*TH*, *DBH*), and sympathoblast markers (*MKI67*, *TOP2A*). Cells are colored based on normalized expression. (D-E) UMAP plots of malignant neuroblastoma data, colored based on adrenergic (D) and mesenchymal (E) signature scores from van Groningen, et al.^20^ (F-G) Violin plot of adrenergic (F) and mesenchymal (G) signature scores^20^ split by developmental state (mesenchymal, adrenergic, and sympathoblast). (H) Heatmap showing inferred transcription factor activity for a curated list of neuroblastoma core regulatory circuit factors^19,61^. A full list of differentially active transcription factors is available in Table S5. Transcription factor activity is colored based on normalized activity. (I) UMAP plot as in (A), colored based on predicted similarity to fetal adrenal medulla cell states from Jansky, et al.^6^ using SingleR^62^. (J) Heatmap of similarity scores between each joint cluster and fetal adrenal medulla cell states. Similarity scores are calculated as normalized Spearman correlations^62^. (K) Overlay comparing snRNA-seq data from an orthotopic patient-derived xenograft (SJNBL012407_X1) to snRNA-seq from the originating patient tumor (HTAPP-194-SMP-251).

The three major neuroblastoma cell clusters had features consistent with the gene expression programs, core regulatory circuits and chromatin landscapes that distinguish ADRN and MES cell lines^19–21^. The sympathoblasts and the eight adrenergic clusters were significantly enriched in an ADRN cell line gene signature^20^ and the three mesenchymal clusters were significantly enriched in the MES cell line signature^20^ (**Figure 3D-G, Table S5**). Moreover, inferred transcription factor (TF) activity (using SCENIC^59,60^) in our mesenchymal cluster (**Figure 3H**; **Table S5**) included multiple TF regulons that had been previously identified in the MES neuroblastoma cell lines^19,20^ including: *ELK4, EGR3, MAFF, IRF1, IRF2, FLI1, RUNX2, FOSL1*, and *FOSL2* (**Figure 3H**; **Table S5**). Conversely, TF regulons in our SYMP and ADRN clusters (*HAND2, PHOX2A, PHOX2B, ISL2*, and *GATA2*) had been previously identified in the ADRN neuroblastoma cell lines and as part of a MYCN core regulatory circuit^19,20,61^ (**Figure 3H**).

### Neuroblastoma cellular heterogeneity reflects a sympathoadrenal developmental hierarchy

To further characterize the 12 malignant cell states in the context of sympathoadrenal development, we compared each profile to a reference atlas of the human fetal adrenal medulla^6^. Consistent with previous reports^2,4,6,8^, most malignant cell/nucleus profiles (82.4%) had highest similarity to ADRN cell types of the sympathoadrenal lineage, representing different stages of sympathetic neuron and chromaffin cell differentiation (**Figure 3I,J** and **S3A,B** and **Table S5**). Another 5.2% of cells/nuclei had highest similarity (SingleR score^62^) to that of proliferating neuroblasts (**Figure 3I,J** and **S3A,B** and **Table S5, Supplemental Information**). Previously, proliferating and postmitotic adrenergic cells were referred to as proliferating neuroblasts and neuroblasts, respectively, but here we use the term sympathoblast (SYMP) to refer to proliferating cells and ADRN to refer to postmitotic cells of the adrenergic lineage (**Figure 2B**).

The mesenchymal cells (8.5%) in our tumors expressed signatures that most closely matched Schwann cell precursors (SCPs) from the fetal adrenal datasets^12^. SCPs are multipotent cells derived from sympathoblasts that can produce both mesenchymal and adrenergic cell types^4,6,12^ (**Figure 2B**). Malignant cells with the highest late SCP score also have high MES scores from prior cell line analyses (**Figure 3E**, and **S3C**), and SOX10, a marker of SCPs in normal fetal adrenal, was identified by SCENIC as active in our MES cells (**Figure 3H**). We confirmed the enrichment of MES clusters with an SCP signature from a second atlas of the human fetal adrenal medulla^8^ (**Figure S3D,E**). We validated that MES cells with the SCP signature are malignant, by comparing profiles from a patient tumor (HTAPP-194-SMP-251) to those from a matched orthotopic patient-derived xenograft^9,63^ (O-PDX) (SJNBL012407_X1): 2.2% and 1.8% of the nuclei profiles in the patient tumor and O-PDX, respectively, had the MES signature (**Figure 3K**). No other non-malignant human cell type was detected within the O-PDX, consistent with our previous observation that only malignant cell types propagate following orthotopic xenotransplantation^9,63^.

Taken together, these data suggest that neuroblastoma tumors have a mixture of proliferating SYMP, differentiating ADRN and MES cells with subsets spanning the different steps of their developmental hierarchy (**Figure 3I,J**). The MES cells express SCP genes as well as genes found in other mesenchymal cell populations (fibroblasts, myofibroblasts, smooth muscle and melanocytes). This is consistent with the fact that MES (S-type) neuroblastoma cell lines have molecular features of Schwann cells, melanocytes, ectomesenchymal derivative and smooth muscle^14^. Fifteen of the tumors in our cohort have very few (<1% of profiled malignant cells), if any, MES neuroblastoma tumor cells (**Figure S3F**), explaining why previous studies with smaller cohorts of patients have not identified MES neuroblastoma cells with the SCP gene expression signature^2,4,6,8^. Importantly, our results cannot be used to definitively infer a cell of origin for neuroblastoma, because there are reports that neuroblastoma cell lines and xenografts can interconvert between MES and ADRN^21,51,64,65^. Instead, our results are consistent with previous work showing that neuroblastomas can recapitulate varying aspects of the sympathoadrenal developmental plasticity^2,4,6,8^, and we have identified interpatient variability in the degree of heterogeneity.

### Differences in neuroblastoma malignant cell composition correlates with outcome

We next tested whether the variation in malignant cell subset composition is associated with clinical features. We found a statistically significant increase in SYMP cell proportions in MYCN amplified tumors, infant cases (< 18 months of age), and untreated cases (**Figure S4A-D**). Conversely, while some (15/55) tumors had very few (<1%) MES neuroblastoma tumor cells (**Figure S3F)**, the differences in MES cell proportions (continuously or discretely) did not significantly associate (FDR>0.05, scCODA) with MYCN amplification, risk group, age, or treatment status (**Figure S4A-D**).

Because different cell states within neuroblastoma are expected to have distinct epigenetic profiles, we reasoned that methylation profiling of tumors could be used to accurately group tumors with different cellular compositions. Indeed, methylation grouping of pediatric cancers have been successful in identifying epigenetic subgroups of brain tumors and sarcoma^66–68^. We combined data from 190 neuroblastoma samples from the National Cancer Institute’s Therapeutically Applicable Research to Generate Effective Therapeutics (NCI-TARGET) initiative with 34 methylation profiles from our cohort (**Figure 1** and **Table S1**). Consensus clustering^69^ partitioned the tumors to four methylation groups, each with samples from both NCI-TARGET and our cohort (**Figure 4A**). Each methylation group was associated with a different combination of clinical and molecular factors (**Figure 4B**). For example, methylation group III was enriched for Children’s Oncology Group low- or intermediate-risk tumors and younger patients (< 18 months), and group IV tumors were enriched for older patients (>18 months) with *MYCN* amplified tumors. Nevertheless, some tumors with *MYCN* amplification or age younger than 18 months were in group II (n=9 and 16, respectively), reflecting the complex underlying biology of neuroblastoma. Next, based on sc/snRNA-seq data in our cohort, group I and IV tumors had a statistically credible increase in the proportion of MES and SYMP tumor cells, respectively (**Figure 4C** and **S4E**; scCODA Bayesian modeling). We validated these findings using digital cytometry with CIBERSORTx bulk deconvolution^70^ of 124 samples from the TARGET cohort with both RNA-seq and DNA methylation data, with deconvolution parameters optimized by training the model using bulk RNA-seq and sc/snRNA-seq data from our samples (**Figure 4D,E**). Consistent with our cohort, deconvolution of TARGET bulk RNA-Seq (n=124) showed that group I tumors had a significantly higher proportion of MES cells, and group IV tumors had a significantly higher proportion of cycling sympathoblast tumor cells (**Figure 4F** and **S4F**).

**Figure 4.**
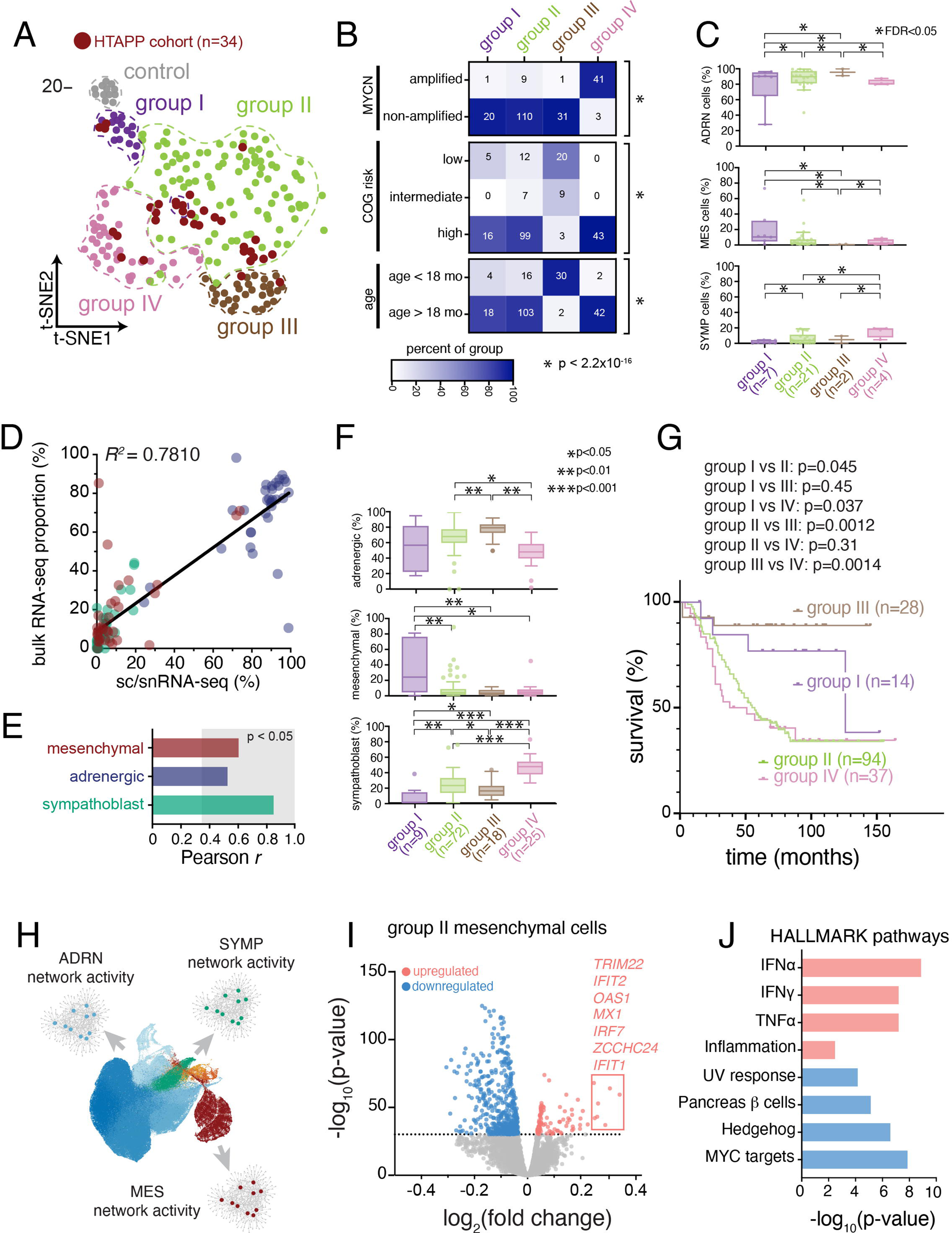
DNA methylation profiling identifies 4 subtypes of neuroblastoma that differ in malignant cell state composition and outcome. (A) t-SNE dimension reduction of 207 neuroblastoma methylation profiles from the NCI TARGET cohort (n=173) or the HTAPP neuroblastoma cohort (n=34). Consensus clustering^69^ was used to delineate 4 groups of tumors plus a group of control adrenal samples. Samples from the HTAPP cohort are demarcated in dark red. (B) Heatmaps colored based on relative abundance of clinical risk factors within each methylation grouping from (A). Numbers within each cell correspond to the absolute number of tumors. p-values, Fisher exact test of independence. (C) Box plot of the proportion of malignant cells/nuclei within each malignant cell state, divided by methylation group. Data are presented as median ± interquartile range. Statistically credible differences, as determined by Bayesian composition analysis^88^ set with a false discovery < 0.05, are displayed as bars with an asterisk. (D and E) Validation and optimization of CIBERSORTx bulk deconvolution parameters derived using matched snRNA-seq and bulk RNA-seq data from n=33 tumors from the HTAPP cohort. (D) shows a scatterplot comparing proportion of cells in each malignant state within sc/snRNA-seq data (‘ground truth’) compared to the estimated proportion from bulk deconvolution. (E) shows the cell-type specific correlation between sc/snRNA-seq and bulk deconvolved data. Bars in the gray region meet significant criteria for concordance with ground truth as measured by Pearson correlation (p < 0.05) (F) Box plots of estimated malignant cell proportions within the NCI TARGET dataset, as estimated by CIBERSORTx bulk deconvolution of bulk RNA-seq data, divided by methylation grouping. Plots show proportions of cells that are estimated to be in the mesenchymal (top), adrenergic (middle), or sympathoblast (bottom) state. Data are presented as median ± interquartile range. Differences across groups that meet statistical significance are shown (p values; Wilcoxon rank sum test). (G) Kaplan-Meier curve showing overall survival of patients within the NCI TARGET cohort (n=175) divided by methylation grouping. p-values were calculated using the Mantel-Cox log rank test. Censored datapoints are represented with solid rectangles. (H) Schematic of the scMINER activity inference. scMINER generates cell type-specific gene regulatory network to infer protein activity and signaling factor activation. (I) Volcano plot of signaling pathway inference comparing group II MES cells to MES cells from the other methylation groupings. (J) HALLMARK pathway analysis of statistically significant results from (I).

Currently, risk stratification of neuroblastoma incorporates multiple clinical, diagnostic, and histologic features to identify patients with a higher risk of death^26,71^. Analyzing the outcome for patients in each of the four tumor groups (**Figure 4G**), we found that patients with group I or III tumors had significantly better outcomes compared to patients with group II or IV tumors (**Figure S4G**). The poor outcomes for group IV patients were consistent with the enrichment of MYCN amplified tumors^26,72,73^ in that group (**Figure 4B**). Similarly, the excellent outcomes for group III patients were consistent with the enrichment of stage 4S patients and infants under 18 months of age^26,27^.Taken together, our data suggest that there may be a correlation between cellular composition, DNA methylation grouping and outcome in neuroblastoma.

As group II tumors had similar proportions of MES and SYMP cells to those of Group III tumors, but had much poorer outcomes, we hypothesized that there may be finer changes in cell states associated with their distinctions. To identify those, we further analyzed cell signaling pathways in each cell population in each disease group. We first used SJARACNe^74^ to infer cell type–specific interactomes for each of the three major cell populations (SYMP, ADRN, MES) from their sc/snRNA-seq profiles, and then used these interactomes to infer the network activity in each nucleus profile using NetBID (**Figure 4H**). Finally, we performed differential activity analysis to identify cell type–specific signaling in each neuroblastoma group (I-IV). Group II MES tumor cells, when compared to MES cells from other methylation groups, were enriched for signatures of inflammatory pathways, including members of the interferon _α/γ_-response and TNF_α_ pathways; in contrast, there were few differences in group II adrenergic cells, when we compared them to adrenergic cells from other methylation groups (**Figure 4I,J, Figure S4H-I**).

### Immune infiltration is correlated with clinical features of neuroblastoma

While bulk methylation profiles identified groups of tumors associated with different clinical outcome, malignant cell composition may be insufficient to fully explain these distinctions (**Figure S4F**). We hypothesized that other cells, such as immune cells, and their interactions with malignant cell states (such as the inflammatory MES state in Group II) may also drive some of these distinctions. To explore this, we characterized the diversity of immune cell states within neuroblastoma. We re-integrated 25,438 cell and 49,515 nucleus profiles from immune cells and annotated them into major categories with CellTypist^75^ (**Figure 5A and S5A-C**). Because tissue dissociation increased stress module gene expression in scRNA-seq (**Figure S5B**), we focused our subsequent analysis on the snRNA-seq profiles.

**Figure 5.**
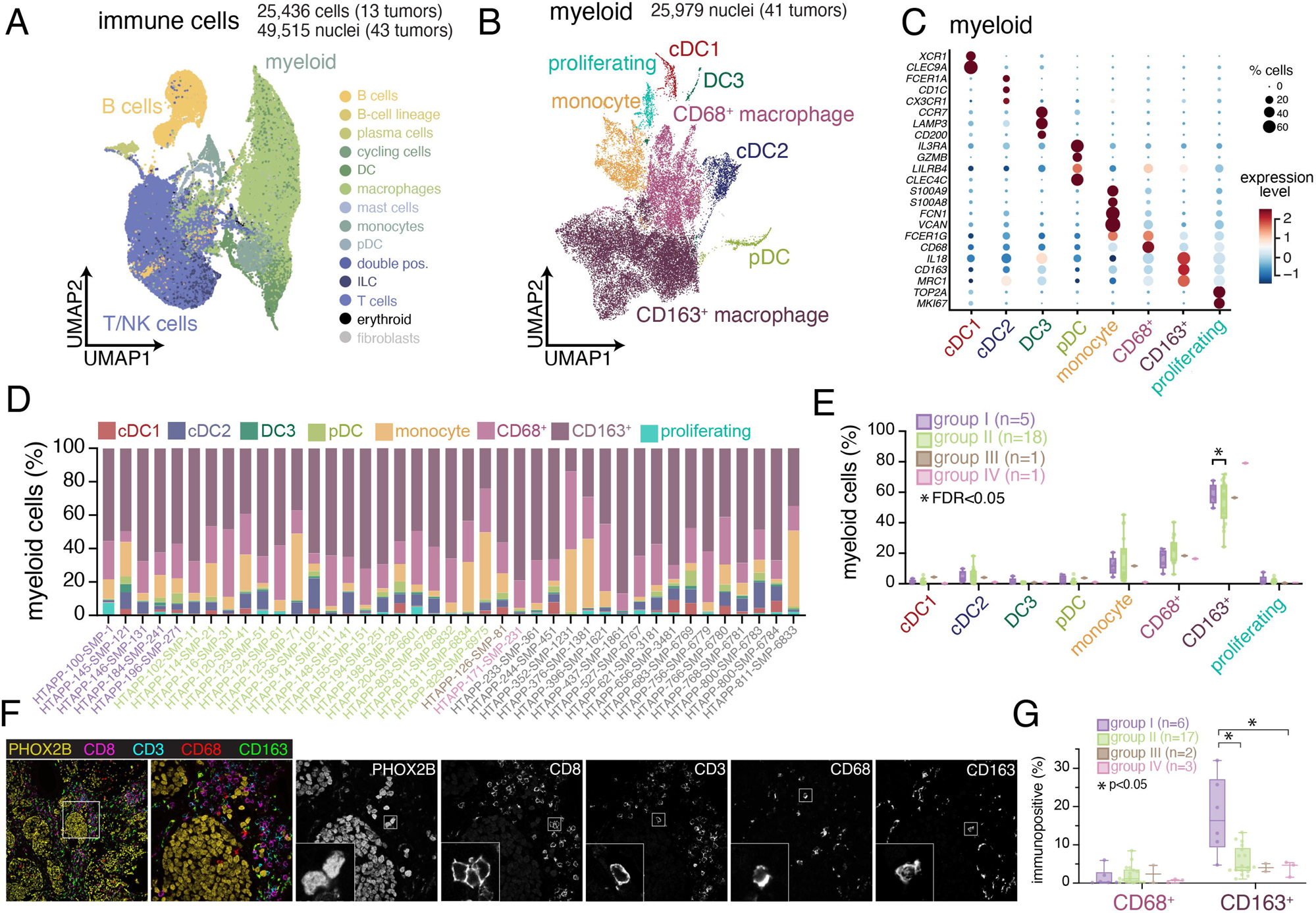
Immune cell heterogeneity and spatial compartmentalization of neuroblastoma. (A) UMAP plot of integrated immune cells/nuclei from scRNA-seq (n=13 tumors; 25,436 cells) and snRNA-seq (n=43 tumors; 49,515 nuclei). Cells/nuclei are colored based on CellTypist automated annotation^75^. (B) UMAP plot of 25,979 myeloid nuclei from the HTAPP neuroblastoma dataset (B). Nuclei are categorized based on the expression of cell markers. (C) Dot plot showing expression of cell markers for each myeloid cluster. Expression is colored based on scaled normalized value (z-score) and the size of each dot represents the percentage of cells within each cluster that had detectable expression. (D) Bar plots showing cell type proportions of myeloid subtypes for each sample within the snRNA-seq cohort (n=41). (E) Composition analysis of the myeloid compartment across snRNA-seq datasets (n=41), separated by methylation grouping. Data are presented as median ± interquartile range. Statistically credible differences, as measured using Bayesian component analysis with false discover rate < 0.05, are represented with an asterisk. (F) Example image of multiplexed immunofluorescence for sample HTAPP-102-SMP-11, stained with PHOX2B, CD8, CD3, CD68, and CD163. (G) Quantitation of CD68+ and CD163+ cells from multiplexed immunofluorescence (n=28). Data is divided by methylation grouping. Data is presented as median ± interquartile range. Differences across groups that meet statistical significance are marked with an asterisk (p values; Wilcoxon rank sum test).

The 25,579 myeloid nucleus profiles from 41 tumors spanned 8 subsets (**Figure 5B-D**). Most myeloid cells (78.9%) were macrophages, including CD68^+^ and CD163^+^ macrophages **(Figure 5B-D)**. CD68^+^ macrophages were enriched in mitochondrial genes (e.g., *MT-CO1/2/3* and *MT-ND1/2/3/4/5*) and genes encoding the ferritin heteromeric complex (*FTL* and *FTH*) (**Table S6**), whereas CD163^+^ macrophages expressed markers of pro-tumorigenic macrophages, including *MDC1* (CD206) and *MSR1* (CD204) (**Table S6**). The 17,142 T/NK cell nucleus profiles included four T cell subsets, one natural killer (NK) cell subset and one small innate lymphoid cell (ILC) subset (**Figure S5D-G** and **Table S6**). Finally, there were 4,962 profiles of naive, memory, and plasma B cells, detected in some, but not all, tumors (**Figure S5H-K** and **Table S6**).

The 25 untreated tumors with snRNA-seq data across methylation Groups I-IV also differed in immune cell compositions (**Figure 5D-G and S5G**). While the number of group III and group IV tumors was limited, there was a statistically credible enrichment of CD163^+^ macrophages in group I *vs*. group II tumors (**Figure 5E**; scCODA Bayesian modeling). We confirmed that that group I tumors had a significantly higher number of CD163^+^ cells compared to tumors from the other methylation groups using 5-plex immunofluorescence microscopy of 28 HTAPP tumors with PHOX2B staining to identify malignant cells and four immune markers (CD3, CD8, CD68 and CD163) to mark T cell and macrophage populations (**Figure 5F,G**). Multiplexed fluorescence showed a compartmentalization phenotype, where PHOX2B^+^ tumor cells were arrayed in nests that excluded immune effector cells and MES cells (**Figure 5F**).

### Discrete spatial compartments of malignant, immune, and stromal cells in neuroblastoma

To better characterize the distinct malignant and immune cell features across neuroblastoma groups, we used multiplex ion beam imaging (MIBI), which combines metal-conjugated antibodies and time-of-flight mass spectrometry^40,41,76^, to generate high-resolution spatial maps of 41 proteins in archival formalin-fixed paraffin-embedded tissue. We first tiled MIBI captures from two high-quality tumor specimens (HTAPP-102-SMP-11 (Group II) and HTAPP-130-SMP-91 (Group IV)) to generate large 2000 μm x 2000 μm arrays (**Figure 6A-C**). We then developed a computational method, Patched Level Analysis of NeuroBlastoma (PLANB), to quantify the expression of markers relative to the tumor and stroma in a stepwise pixel-by-pixel manner (**Figure 6D**). The expression of CD56/NCAM (a marker of neuroblastoma^77^), proliferation (Ki67), and transcriptionally active chromatin (H3K27ac) were all enriched within tumor neighborhoods (**Figure 6D**), whereas markers for immune effectors (CD11c, CD4 and CD8) were enriched in the stromal neighborhood (**Figure 6D**). We next performed MIBI analysis using 30 fields of view (FOVs) from 11 patient samples, predominantly from Group II (n=7). We annotated cell types based on protein expression (**Figure 6E-G**), and observed both sample-to-sample and region-to-region variation in immune cell density that correlated with the sc/snRNA-seq (**Figure 6G,H**). Neighborhood analysis (n=30) identified broad patterns in cell type colocalization (**Figure 6I,J**). Consistent with PLANB, Malignant tumor cells segregated into one neighborhood (neighborhood ‘0’) with very few immune cells (**Figure 6J**, neighborhood ‘0’), while there were four distinct archetypes of immune-rich neighborhoods. For example, neighborhood ‘3’ was particularly rich in B cells, whereas macrophages were predominantly localized to neighborhood ‘2’ that also contained CD4 T, CD8 T, and dendritic cells (DCs) (**Figure 6J,K**).

**Figure 6.**
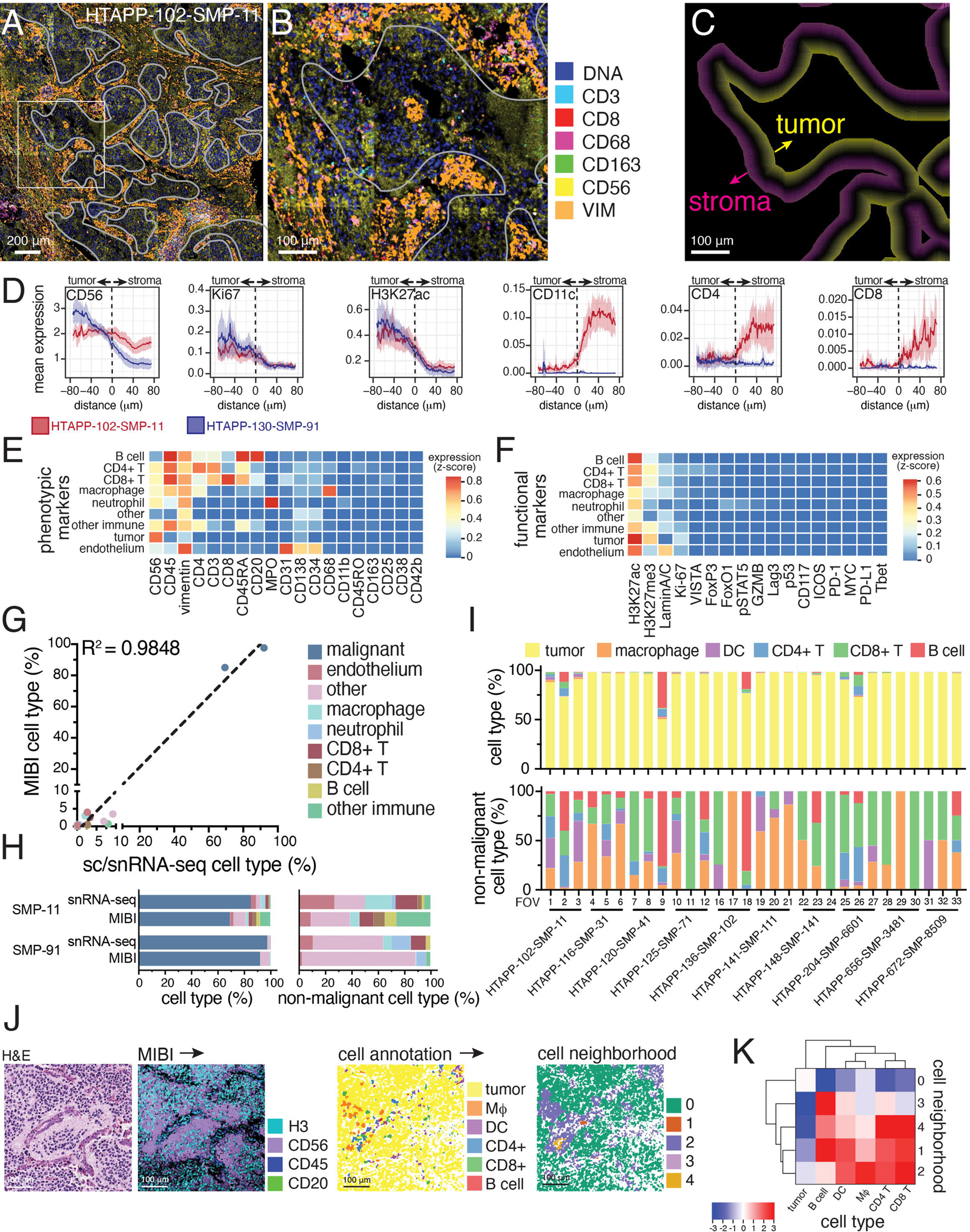
Multiplexed ion-beam imaging (MIBI) identifies compartmentalization of tumor and immune cells. (A) Multiplexed ion beam imaging (MIBI) of multiplexed spatial proteomic data. Two large, tiled arrays (5×5 captured areas) from samples HTAPP-102-SMP-11 and HTAPP-130-SMP-91 were obtained. As an example, the stitched array from HTAPP-102-SMP-11 is shown in panel A; 7 markers are shown (double-stranded DNA [dsDNA], CD3, CD8, CD68, CD163, CD56 and Vimentin [VIM]). A white overlay shows the tumor-stroma interface, which was used for patched level analysis of neuroblastoma (PLANB). (B) A high magnification of the boxed area within (A). (C) Patched level analysis of neuroblastoma (PLANB) determines the relative locality of cell types or markers with reference to the tumor-stroma boundary using a stepwise pixel-by-pixel algorithm. (D) PLANB analysis of HTAPP-102-SMP-11 and HTAPP-130-SMP-91 showing the distribution of tumor markers (CD56, Ki67, H3K27ac), myeloid cells (CD11c), or T cells (CD4, CD8). The x-axis reflects distance relative to tumor-stroma interface, with negative values being within the tumor nest and positive values being outside the tumor nest. The y-axis reflects relative expression of markers (arbitrary units). (E-F) Heatmap showing expression of cell type phenotypic markers (E) and functional markers (F) for each cell type in the large, tiled arrays. (G-H) Cell type proportions from the two large, tiled arrays comparing the prevalence of cell types in the MIBI data compared to snRNA-seq data. Panel (G) shows a scatterplot comparing proportions of each cell type identified in sc/snRNA-seq data (*x*-axis) to the proportion detected in the tiled MIBI arrays for HTAPP-102-SMP-11 and HTAPP-130-SMP-91. Panel (H) shows bar plots comparing cell type composition in snRNA-seq and MIBI datasets both including (*left*) and excluding (*right*) malignant cells/nuclei. (I) Bar plot of cell types in the single-tile MIBI datasets (10 samples, 30 FOVs). Following capture of MIBI data, segmentation and marker expression was used to annotate cell types. Bar plots including and excluding malignant cells are shown on the top and bottom, respectively. (J) Neighborhood analysis of MIBI data from the 30 FOVs from (H). As an example, we show HTAPP-102-SMP-11 field-of-view (FOV) #1. On the left side, we show an H&E from an adjacent section. Following MIBI capture, images were segmented, and cell types were automatically called. These cell maps were then analyzed using neighborhood analysis to identify those cell types that co-localize with each other. (K) Cluster-map of cell type composition within each neighborhood from the combined MIBI dataset (n=30 FOVs). Dendrograms represent hierarchical clustering of cell types (rows) and neighborhoods (columns). Colors represent scaled likelihood of adjacency between cell types.

Because MIBI provided a high-resolution map of neuroblastoma spatial organization but could not be performed at scale for large areas across the samples, we generated whole-slide CODEX^44,78^ data for 10 HTAPP samples (with matching MIBI data) with 52 antibodies followed by imaging segmentation, cell marker clustering, and cell annotation (**Figure 7A-C** and **S6A-C**). We leveraged the expanded scale of CODEX to characterize the inter- and intra-tumor cellular heterogeneity across the entire tumor section for each sample. First, we partitioned CD56^+^ CD45^−^ cells by expression of vimentin (VIM), collagen IV (COL4), and podoplanin (PDPN). CD56^+^VIM^+^COL4^+^PDPN^+^ cells had thin, elongated morphology and were adjacent to neuropil, consistent with Schwannian stroma (**Figure S6D-F**). CD56^+^VIM^−^COL4^−^PDPN^−^ cells had histologic features of neuroblastoma tumor cells (small blue round cells) and were positive for the proliferation marker Ki67 (**Figure S6D-F** and data not shown). CD56^+^VIM^+^COL4^+^PDPN^−^ and CD56^+^VIM^+^COL4^−^PDPN^−^cells were rare and interspersed throughout the tumor (**Figure 7C** and **S6F**). We cannot discern from these data if those cells are malignant. Overall, because of the available antibodies, CODEX was more useful for spatial analysis of immune than neuroblastoma malignant cell types (ADRN, MES, SYMP).

**Figure 7.**
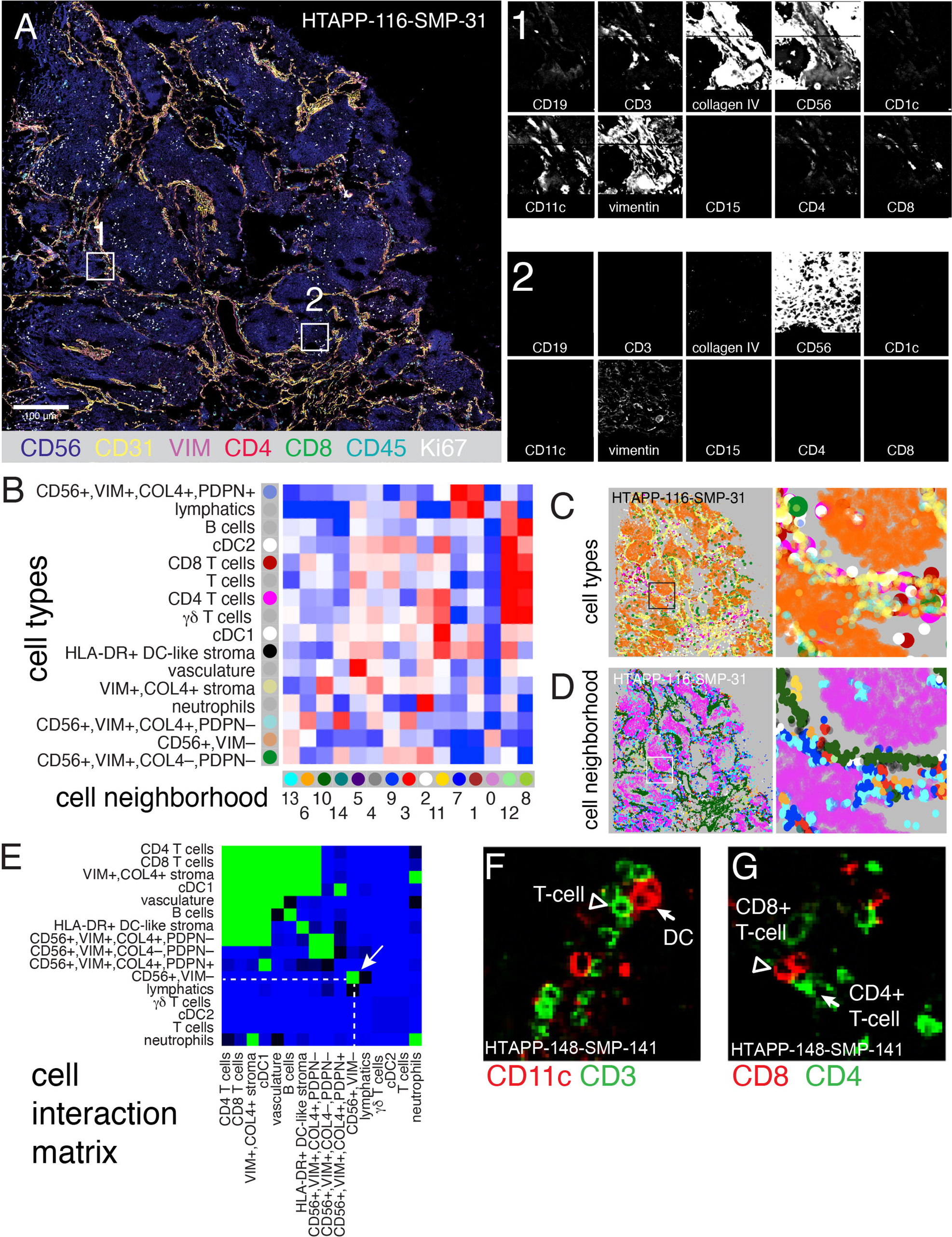
Co-detection by indexing (CODEX) imaging of whole slides identifies recurrent patterns of cellular interaction. (A) Representative CODEX whole-slide image of HTAPP-161-SMP-31, showing 3-color overview (using CD56, collagen IV [COLIV] and CD3), imaged using a 52-marker CODEX panel. Insets of two regions are shown with a selected panel of markers typical for B cells (CD19), T cells (CD3, CD4, and CD8), stroma (collagen IV), tumor (CD56), myeloid cells (CD1c, CD11c, and CD15), and cells of ambiguous lineage (Vimentin). (B) Identification of 15 cell neighborhoods (CNs) based on 16 cell types, showing cell type enrichment within each CN (pooled data across 10 CODEX samples). (C) Representative cell type maps from sample HTAPP-116-SMP-31. Cells are colored to match the row legend in panel B. (D) Representative CN map from sample HTAPP-116-SMP-31. Neighborhoods are colored to match the column legend in panel B. (E) Heatmap of likelihood ratios of cell-cell contact between the 16 annotated cell types across the entire HTAPP CODEX cohort (n=10 samples). (F-G) Representative CODEX images of cell contacts between either CD3-positive T cells and CD11c-positive dendritic cells (F) or CD4-positive and CD8-positive T cells (G).

Following cell type annotation, we identified cell neighborhoods (CNs) with characteristic co-localized cell types (**Figure 7B-D**). The malignant CD56^+^ cells were predominantly found in CN0, which contained no other cell types (**Figure 7B,D**), consistent with our MIBI and multiplex immunofluorescence analysis. Three other neighborhoods (CN8, CN11, and CN12) were predominantly enriched for immune cell types, especially DCs and T cells (**Figure 7B,D**). Finally, seven neighborhoods were predominantly comprised of stromal cells, including two lymphatic-rich neighborhoods (CN1 and CN7) and five cell neighborhoods that were enriched for VIM+,COL4+ stroma (CN2, CN3, CD4, CD5, and CN10; **Figure 7B,D**).

Unsupervised analysis of the cell-cell interaction matrix from the entire tumor section of each of the 10 samples confirmed the compartmentalization of tumor cells from immune/stromal components in neuroblastoma (**Figure 7E-G**). Specifically, CD56^+^ tumor cells self-associate into a niche devoid of immune cells (**Figure 7E**). Each tumor cells neighborhood is in turn surrounded by a second niche with stromal cells, dendritic cells, and T cells (**Figure 7E-G**). A third niche comprised of CD56^+^VIM^+^ cells that may represent tumor cells, normal cells, or both. Taken together, both CODEX and MIBI showed a highly compartmentalized tumor organization.

### High-resolution spatial transcriptomics shows context-dependent shifts in expression

As the number and type of antibodies used in spatial proteomics limited our ability to resolve malignant cells, we finally profiled 19 frozen sections from 10 tumors in our cohort by Slide-SeqV2^42^, generating spatial transcriptomic data at 10 μm resolution. We then integrated our Slide-SeqV2 and snRNA-seq profiles to spatially assign cell type distributions (with robust cell type decomposition (RCTD)^79^), overcoming the spatial lower resolution, and to spatially project full gene expression profiles (using TANGRAM^80^), overcoming data sparsity in Slide-SeqV2 (**Figure 8A**).

**Figure 8.**
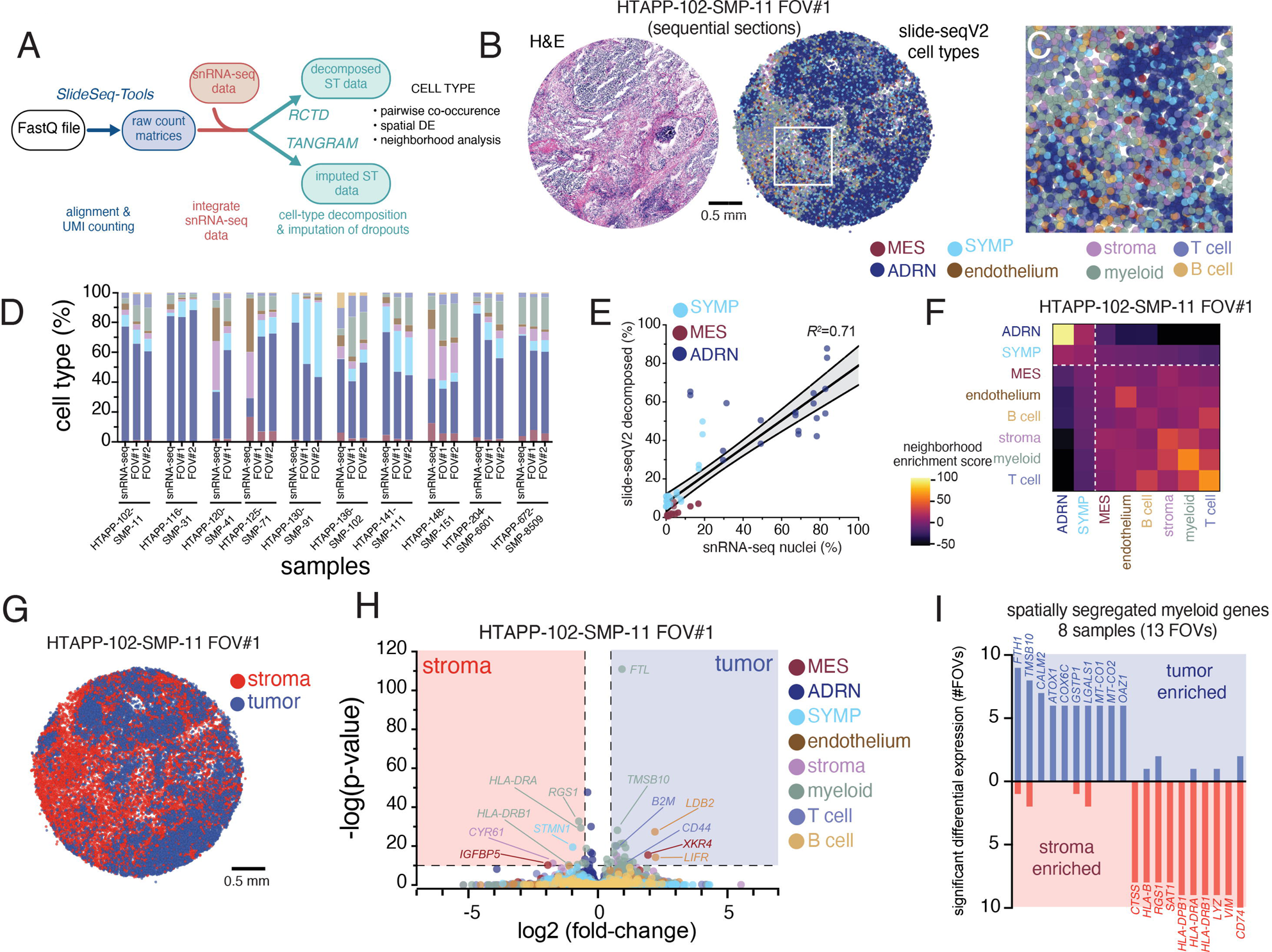
High resolution spatial transcriptomics with Slide-SeqV2 demonstrates myeloid reprogramming within neuroblastoma. (A) Data processing workflow for Slide-Seq v2 data. Slide-SeqV2, after alignment and UMI counting, underwent both cell type decomposition and zero-count imputation. For both analyses, analogous snRNA-seq from the same tumor fragment was employed to define cell types or expression modules. (B) An example Slide-SeqV2 array from HTAPP-102-SMP-11. On the left, an adjacent frozen section was processed by H&E staining. On the right, a Slide-SeqV2 array colored based on cell type after decomposition with RCTD^79^. (C) Magnified view of the HTAPP-102-SMP-11 field-of-view (FOV) #1 from panel B. (D) Comparison of cell composition from snRNA-seq data and Slide-SeqV2 assays. 17 Slide-SeqV2 datasets were generated from 10 specimens. The colors of the bars match up with the legend in panels B and C. (E) Scatterplot comparing the percentage of malignant cell states within single-nucleus RNA-seq data (x-axis; n=10) to the percentage of measured within each decomposed Slide-SeqV2 FOV (y-axis; n=19). (F) Cell-cell interaction cluster-map of cell types showing the frequency that two cell types are adjacent to each other. (G) Tumor-rich and stroma-rich compartments, defined after clustering. Tumor-rich areas were defined as Slide-SeqV2 beads with >50% malignant cell composition while stroma-rich areas were defined as Slide-SeqV2 beads with <50% malignant cell composition. (H) Volcano plot of CSIDE differential expression analysis comparing cell-type expression patterns between tumor-rich and stroma-rich compartments. (I) Bar plot of recurrent myeloid genes that are enriched in tumor-rich myeloid cells or stromal-rich myeloid cells. The y-axis represents the frequency that significant enrichment was detected (out of 13 evaluated FOVs).

We successfully assigned 8 of the cell types defined in sc/snRNA-seq (**Figure 8B-D**), including all three major malignant cell subsets (ADRN, MES, SYMP), three broad immune cell categories (T cells, B cells, myeloid cells), and stromal and vascular endothelial cells. The proportion of malignant cell populations was correlated (Pearson’s r^2^=0.71, p<0.0001) between snRNA-seq and Slide-SeqV2 (**Figure 8E**). We then used the relative abundance of cell signatures in Slide-SeqV2 data to identify cell-cell colocalization, finding that ADRN and SYMP tumor cells were less likely to be adjacent to myeloid or T-cells (**Figure 8F**), consistent with the compartmentalization observed in spatial proteomics analysis (MIBI and CODEX). In contrast, MES tumor cells had a weak, but detectable, association with immune cells in the TME (**Figure 8F**), which is consistent with the higher expression of an inflammatory program by MES cells in Group II tumors (**Figure 4I,J**).

To investigate further, we clustered the RNA profiles from each barcoded bead and then aggregated those clusters into a tumor-rich (>50% malignant cells) and stroma-rich (<50% malignant cells) beads (**Figure 8G**), and then compared for each cell type, differential expression (per section) between beads with that cell type that are in the tumor-rich *vs*. stroma-rich clusters (**Figure 8H-I**). For example, myeloid cells in sample HTAPP-102-SMP-11 FOV#1 expressed significantly higher ferritin light chain (*FTL*) when in tumor-rich *vs*. stroma-rich beads (**Figure 8H**), and higher levels of MHC class II genes *HLA-DRA* and *HLA-DRB1* in stroma-*vs*. tumor-rich beads (**Figure 8H**). Finally, we scored genes for recurrent enrichment in either compartment across the 19 samples (**Figure 8I** and **Table S7**). Stroma-restricted myeloid cells consistently expressed monocytic and DC markers including MHC class II genes (*HLA-DPB1*, *HLA-DRA*, *HLA-DRB1*, *CD74*) and lysozyme (*LYZ*) (**Figure 8I**), whereas intratumoral myeloid cells expressed higher levels of mitochondrial (e.g., *MT-CO1* and *MT-CO2*) and anti-oxidant (e.g., *ATOX1*, *GSTP1*, *COX6C*, and *OAZ1*) genes, raising the possibility that those myeloid cells that penetrated into tumor-rich regions underwent metabolic stress (**Figure 8I**). Indeed, the CD68^+^ macrophages snRNA-seq profiles (**Figure 5B,C**) are enriched for *FTL* and *TMSB10*, both associated with intratumoral expression in Slide-SeqV2, supporting this interpretation.

## DISCUSSION

Tumor profiling using single-cell sequencing technologies has transformed our understanding of cancer and the cellular constituents within tumors. Spatial-omics technologies have the potential to unveil further complexity informed by the tissue structure within tumors. Despite these advances, combining and leveraging these tools in concert remains limited due to technical and computational challenges. We have generated a comprehensive dataset for a rare childhood cancer, neuroblastoma, by accruing fresh and frozen samples from two pediatric cancer institutions. We have developed standardized tissue-processing and handling methods to decentralize data generation, which we have made publicly available to the community^45^ (Methods). In total, we used four sequencing methods (single-cell/nucleus RNA-seq, bulk RNA-seq, whole exome sequencing and methylation profiling) and three spatial methods (MIBI, CODEX and Slide-SeqV2) to build a foundational resource for pediatric cancer researchers and for future computational method development. We have made all data (raw, processed and annotated) openly available to the research community through the HTAN Data Coordinating Center (https://humantumoratlas.org/). In this study, we leverage this resource to investigate heterogeneity of malignant cell states and to map the spatial organization of immune cells within neuroblastoma.

### Malignant cells in neuroblastoma retain features of sympathoadrenal development

Our analysis of single cell/nucleus RNA-sequencing profiles from 55 samples show that neuroblastoma tumors have three major cell populations that are reminiscent of sympathoadrenal development. Most malignant cells resemble differentiating adrenergic neurons (post-ganglionic sympathetic neurons and chromaffin cells; ADRN cells). A smaller subset of cells resembles the immature proliferating sympathoblasts from early stages of sympathoadrenal development (SYMP). A diverse nomenclature has been used to describe these cells, which have been referred to as ‘sympathoblasts’, ‘neuroblasts’, ‘adrenergic cells’, and ‘noradrenergic neuroblastoma’ ^6,8,14,19,20^. We propose restricting the term sympathoblast to those cells that are actively dividing in sc/snRNA-seq datasets, and the term adrenergic for all other cell populations that have features of sympathoadrenal neurons and chromaffin cells. In the future, the community may define further subsets of cells in the ADRN population of neuroblastoma tumor cells that resemble different stages of development.

In addition to ADRN and SYMP tumor cells, we identified a third ‘mesenchymal’ (MES cells) population of tumor cells. The presence of these mesenchymal cells hearkens to previous reports using cell lines, a subset of which were shown to have epigenetically and transcriptionally similarity to early neural crest progenitors^19,20^. In our single-cell/nucleus dataset, the MES cells/nuclei had expression profiles consistent with those reports providing further evidence of this important biological insight^19,20^. However, other neuroblastoma single-cell studies have raised the question of whether MES cells are bona fide malignant cells. Indeed, a variety of non-malignant cells (Schwann cells, SCPs, fibroblasts, myofibroblasts) that express mesenchymal genes are present in neuroblastoma tumors, making it difficult to discern bona fide MES neuroblastoma cells from non-malignant cells expressing similar gene expression programs^81^. In our analysis, we separated MES and ADRN malignant cells and showed that both populations had the same inferred copy number variation. Additionally, using datasets from matched patient tumor and an orthotopic patient-derived xenograft (O-PDX), we show that SYMP, ADRN and MES cell populations are present in similar proportions. Non-malignant human cells do not persist during passaging in O-PDXs, lending further credence that the MES tumor cell population are indeed malignant cells. Deeper analysis of the MES neuroblastoma cell population showed these cells most closely resembles Schwann cell precursors (SCPs) from the developing fetal adrenal medulla, consistent with a recent report that also identified an SCP-like population in a cohort of 17 tumors^36^. While the similarity of MES and SCP expression profiles raise the tantalizing possibility that neuroblastoma recapitulate the differentiation hierarchy of the developing fetal adrenal, it remains unclear whether SCPs are the cell of origin of neuroblastoma and future work will be needed to address that question.

### DNA methylation correlates with clinical outcome and tumor heterogeneity

We observed a broad range of heterogeneity patterns, including 15 neuroblastomas samples which were deficient of MES cells (defined by having less than 1% of malignant cells/nuclei in the MES subpopulation). Traditional clinical risk factors such as age, risk strata, and MYCN amplification status did not credibly distinguish the degree of tumor cell heterogeneity. DNA methylation, however, identified four molecular groups with distinct patterns of malignant state heterogeneity. These groupings are reminiscent of a recent study by Gartlgruber, et al. that used histone H3K27ac profiling to identify four epigenetic groups of neuroblastoma^82^. They discovered one group of tumors enriched for MYCN amplified tumors, and two groups that had MYCN non-amplified tumors differing in risk status; interestingly, a fourth group that the authors reported as “mesenchymal subtype” shared transcriptomic similarity to SCPs. In our analysis, we were able to take advantage of matched methylation and transcriptomic data (both bulk RNA and single-cell/nucleus RNA-sequencing) to directly compare the malignant cell state diversity within each methylation group. Indeed, we also identified a group of tumors (group I) that were highly enriched for MES tumor cells. In group II tumors, MES cells expressed higher levels of pro-inflammatory gene programs indicative of interferon and TNF pathway activation, consistent with in vitro reports that cells with a MES transcriptomic signatures had higher levels of MHC class II expression and were able to engage immune effector cells^83,84^. Group III tumors were enriched for young infants, and most group IV tumors had MYCN amplifications. Importantly, though each methylation group had distinct patterns of clinical risk factors, they were a number of outliers. For example, we identified 9 MYCN amplified tumors that clustered into group II, and 2 infants clustered into group IV.

Critically, methylation grouping correlated with survival outcomes, which raises the possibility of using DNA methylation to molecularly stratify neuroblastoma risk. DNA methylation profiling has already shown utility in the diagnostic classification of brain tumors and sarcomas^66,67^ and has been validated in the risk stratification of medulloblastoma^85^. Before any similar implementation in neuroblastoma, much larger methylation datasets will need to be gathered to construct a definitive epigenetic classifier. Finally, it remains unclear whether transcriptomic or epigenetic shifts during therapy could be used as a prognostic biomarker. Prior studies in preclinical models of neuroblastoma have indicated that relapsed neuroblastoma tumors are enriched for mesenchymal transcriptomic signature^20,65^, and transcription factor activity analysis of 3 matched primary-relapsed samples indicated an association between the MES signature and disease relapse^82^. Future work comparing transcriptomic or methylation profiles from tumors samples obtained before and during therapy may provide valuable information about mechanisms of treatment escape and recurrence in neuroblastoma.

### Spatial-Omics

Spatial profiling technologies have dramatically improved our ability to map the cellular architecture of tumors in situ. However, their application to rare childhood cancers has been limited by the lack of a standardized dataset comparing the strengths and limitations of each modality. By generating MIBI, CODEX and Slide-SeqV2 data from 10 banked neuroblastoma tumors, we benchmark each assay so that they compared side-by-side. MIBI provides very high spatial resolution which is useful for analysis of direct cellular interactions in tumors and the TME. Another advantage is the use of archival FFPE sections with excellent preservation of morphological features. The limitation of MIBI is the size of the fields that can be efficiently analyzed. We produced large, tiled datasets for two tumors, but had to use smaller fields of view for the remainder of our cohort. CODEX overcomes this limitation, and we were able to generate whole slide images; at the time that we were acquiring data, CODEX had been credentialed only for use with fresh-frozen tissue, but interim advances have adapted CODEX for use with FFPE tissue. Both MIBI and CODEX are limited by the availability of antibodies that have been validated in the tissue of interest, and the majority of validated antibodies target immune cell markers. Thus, these two platforms were particularly useful for profiling the immune cells in the TME, but the antibodies available could not be used to definitively identify all 3 tumor cell populations. Slide-SeqV2, however, provided broader data on cell populations in the tumor and TME, allowing us to identify patterns of spatial distribution for each malignant cell state. Slide-SeqV2 had the limitation of reduced resolution (10 μm), which required integration with sc/snRNA-seq data to map cell types across the specimen. Nonetheless, if there are well-separated regions of the tumor and TME that can be defined histologically, Slide-SeqV2 can be useful in identifying differences in gene expression patterns for the same cell population across the different neighborhoods in the tumor. We envision, that based on our experience, that future atlas efforts using a combination of a spatial proteomic, spatial transcriptomic, and single-cell RNA-sequencing technologies will be able to generate the greatest cross-modality information. Further advances in computational tools that allow researchers to bridge between spatial -omic data will be needed to fully harness these multi-modal atlases; indeed two recent reports have shown the promise of this approach by unifying cell type annotation^86,87^.

Taken together, all three platforms were consistent with the conclusion that neuroblastomas have discrete tumor cell neighborhoods made up of SYMP and ADRN tumor cells surrounded by a stromal neighborhood made up of MES tumor cells, immune cells, and other cells in the TME. This is important because the assumption that neuroblastomas are immunologically ‘cold’ may reflect this segregation of cell populations rather than a complete lack of immune cells in the tumor. Moreover, it suggests that different populations of tumor cells may play different roles in signaling to the immune cells in the TME. For example, our data indicate that MES cells within the TME have activation of pro-inflammatory genes and pathways, and this may contribute to the compartmentalization of neuroblastoma. Also, these platforms were useful in comparing the cell-cell signaling and gene expression network activity in the cell populations that were in different regions of the tumor such as macrophages in the tumor neighborhood relative to the stromal regions. We have demonstrated the importance of integrating multiple spatial-omics platforms with sc/snRNA-seq and other molecular and cellular features to gain a more comprehensive view of the cellular heterogeneity and interactions in cancer.

### Future Directions

To provide clarity for the neuroblastoma field, it will be important to harmonize the nomenclature of cell states relative to normal fetal adrenal development. We propose the use of the term ‘SYMP’ for the proliferating sympathoblast cells and ‘ADRN’ for the more differentiated tumor cells that are not actively dividing. This is consistent with the previous studies and the historical research on cell lines. Along those lines, we propose the term ‘MES’ to refer to the non-neuronal cells in neuroblastoma that have mixtures of gene expression programs reminiscent of mesenchymal cell populations derived from SCPs. In addition to simplifying and clarifying the nomenclature, it will be essential to independently validate the presence of all three cell populations in patient tumors and O-PDXs and to confirm that these three populations vary across patients and correlate with outcome and DNA methylation group. Finally, it will be important to extend our discovery beyond the samples here that neuroblastomas are partitioned into neighborhoods with ADRN and SYMP cells that are separated from the stroma with MES tumor cells and immune cells.

There are exciting opportunities to begin to explore the changes in gene expression programs as cells (e.g., macrophages) migrate between the different neighborhoods within tumors and the changes that occur in the context of treatment. Also, studies on patient tumors undergoing anti-GD2 immunotherapy can provide important new insights into the molecular and cellular mechanisms of antigen-directed cell-mediated cytotoxicity, in order to help us better understand why some patient tumors respond better than others. Finally, there are opportunities to further refine cell populations such as the eight ADRN groups in neuroblastoma and the lineage relationships between tumor cell populations. We do not know if individual tumor cells can transition between the SYMP, ADRN and MES cell states or if they are clonally restricted. Nor do we know if one population of tumor cells is more likely to survive therapy and contribute to disease recurrence. The data generated from this study provide a systematic framework for the future investigation of neuroblastoma. As such, we have shared all data, raw and processed, publicly available through the Human Tumor Atlas Network data portal (https://humantumoratlas.org/). We anticipate that this cohort will be a transformative resource for further dissection of neuroblastoma biology and for building computational tools in the future.

## ACKNOWLEDGEMENTS

Several figure diagrams (Figure 2B, 8A, and S1A) were created with Biorender.com. We thank the St. Jude Children’s Research Hospital Hartwell Center for Bioinformatics for sequencing support. We thank Anna Hupalowska and Chris Fiveash for graphical illustration. We thank the St. Jude Comparative Pathology Core laboratory for immunohistochemistry support.

This project has been funded in part with federal funds from the NCI, National Institutes of Health, task order no. HHSN261100039 under contract no. HHSN261201500003I. The content of this publication does not necessarily reflect the views or policies of the Department of Health and Human Services, nor does mention of trade names, commercial products or organizations imply endorsement by the US Government. B.E.J. received funding through U2C CA233195 and contract 17X149Q2. Additionally, this work was supported by an NCI Cancer Center Support Grant (CA21765). M.A.D. received funding support through the NIH grants EY30180, CA219686, CA248432, and CA24550. The content is solely the responsibility of the authors and does not necessarily represent the official views of the National Institutes of Health.

M.A.D. and A.Regev were supported by the Howard Hughes Medical Institute. M.A.D. was supported by the Tully Family and Peterson Foundations. A.G.P. received funding through Hyundai Hope on Wheels Foundation and Damon Runyon Cancer Foundation (#DRSG-33P-20). Both M.A.D. and A.G.P. received funding from Alex’s Lemonade Stand Foundation. Research was also supported by American Lebanese Syrian Associated Charities.

Lastly, we would like to acknowledge our patients and their families, without whom this work would not have been possible.

## AUTHOR CONTRIBUTIONS

Conceptualization: M.R.C., B.E.J., O.R-R., A.Regev, M.A.D.

Data curation: A.G.P., O.A., N.B.C., S.J., B.A.O., C.Cline, Y.G.

Formal analysis: A.G.P., O.A., N.B.C., S.J., X.H., H.S., C.B.M.P., Y.B., B.Z., Y.G.

Funding acquisition: G.P.N., O.R-R., A.Regev, M.A.D.

Investigation: A.G.P., O.A., N.B.C., Å.S., S.J., M.S., C.Caraccio, H.S., R.G.T., M.L., D.D.,

C.B.M.P., J.Waldman, Y.B., B.Z., I.B., E.M., S.V., S.N., I.W., J.Wu, G.G., A.L., N.C., A.T., S.R., Y.G.

Methodology: A.G.P., O.A., N.B.C., Å.S., S.J., M.S., D.D., C.B.M.P., Y.B., B.Z., S.V., E.S., F.C., Y.G.

Project administration: A.G.P., O.A., N.B.C., Å.S., S.J., C.Cline, L.D., K.H., A.Rotem, K.L.P.,

Å.K., J.J-V., E.T., B.E.J., Y.G., G.P.N., O.R-R., A.Regev, M.A.D.

Resources: A.G.P., O.A., N.B.C., Å.S., S.J., B.A.O., J.D., B.G., E.S.

Software: A.G.P., O.A., S.J., X.H., H.J., K.X., T-C.C., A.Shirinifard, R.H.C., A.Shen, C.B.M.P., Y.B., B.Z., P.G., G.W., J.Y., Y.G.

Supervision: A.G.P., O.A., N.B.C., Å.S., S.J., M.R.C., S.R., B.E.J., F.C., J.Y., Y.G., G.P.N.,

O.R-R., A.Regev, M.A.D.

Validation: A.G.P., O.A., Å.S., S.J., M.S., H.S., Y.B., B.Z., Y.G.

Visualization: S.J., R.H.C., C.R., J.P., P.G., X.Z., Y.G.

Writing - original draft: A.G.P., O.A., N.B.C., Å.S., M.A.D.

Writing - reviewing & editing: A.G.P., O.A., N.B.C., Å.S., B.E.J., A.Regev, M.A.D.

