## Supplemental Information for "A spatial cell atlas of neuroblastoma reveals developmental, epigenetic and spatial axis of tumor heterogeneity"

**Supplementary Information**

**SUPPLEMENTAL TABLES**

**Table S1. Clinical characteristics of neuroblastoma samples used in this study, related to Figures 1.**

**Table S2. Whole exome sequencing data from the HTAPP neuroblastoma cohort, related to Figure 1.**

**Table S3. Single cell/nucleus RNA-seq quality control metrics, related to Figures 2 and 3.**

**Table S4. Gene expression within the combined HTAPP sc/snRNA-seq dataset, related to Figure 2.**

**Table S5. Gene expression within the malignant cells/nuclei from sc/snRNA-seq, related to Figure 3.**

**Table S6. Gene expression within the T cell, myeloid and B cell nuclei, related to Figure 5.**

**Table S7. CSIDE analysis of Slide-SeqV2 data to identify cell-type specific DE genes, related to Figure 8.**

**Table S8. Antibodies used for spatial proteomic analysis, related to STAR Methods.**

**SUPPLEMENTAL FIGURES**

**Figure S1. Workflow and technical validation of sc/snRNA-seq data, related to Figure 2.**

(A) Schematic of the analytic pipeline for single-cell/nucleus RNA-seq within this study.

(B) UMAP plot of all sc/snRNA-seq data highlighting those data points which were obtained from single-cell data (left) or single-nucleus data (right).

(C) UMAP plots of the neuroblastoma transcriptomic atlas, colored based on patient sample.

(D) Montage of UMAP plots for each individual sample within the HTAPP cohort. Single-cell RNA-sequencing datasets are labelled in blue, single-nucleus RNA-sequencing datasets are labelled in brown.

(E-H) Quality control metrics of data from the neuroblastoma transcriptomic atlas comparing genes/cell (E), mitochondrial (mt) reads (F), and dissociation metrics (G,H) from scRNA-seq and snRNA-seq data. p-values calculated using the Wilcoxon signed rank test.

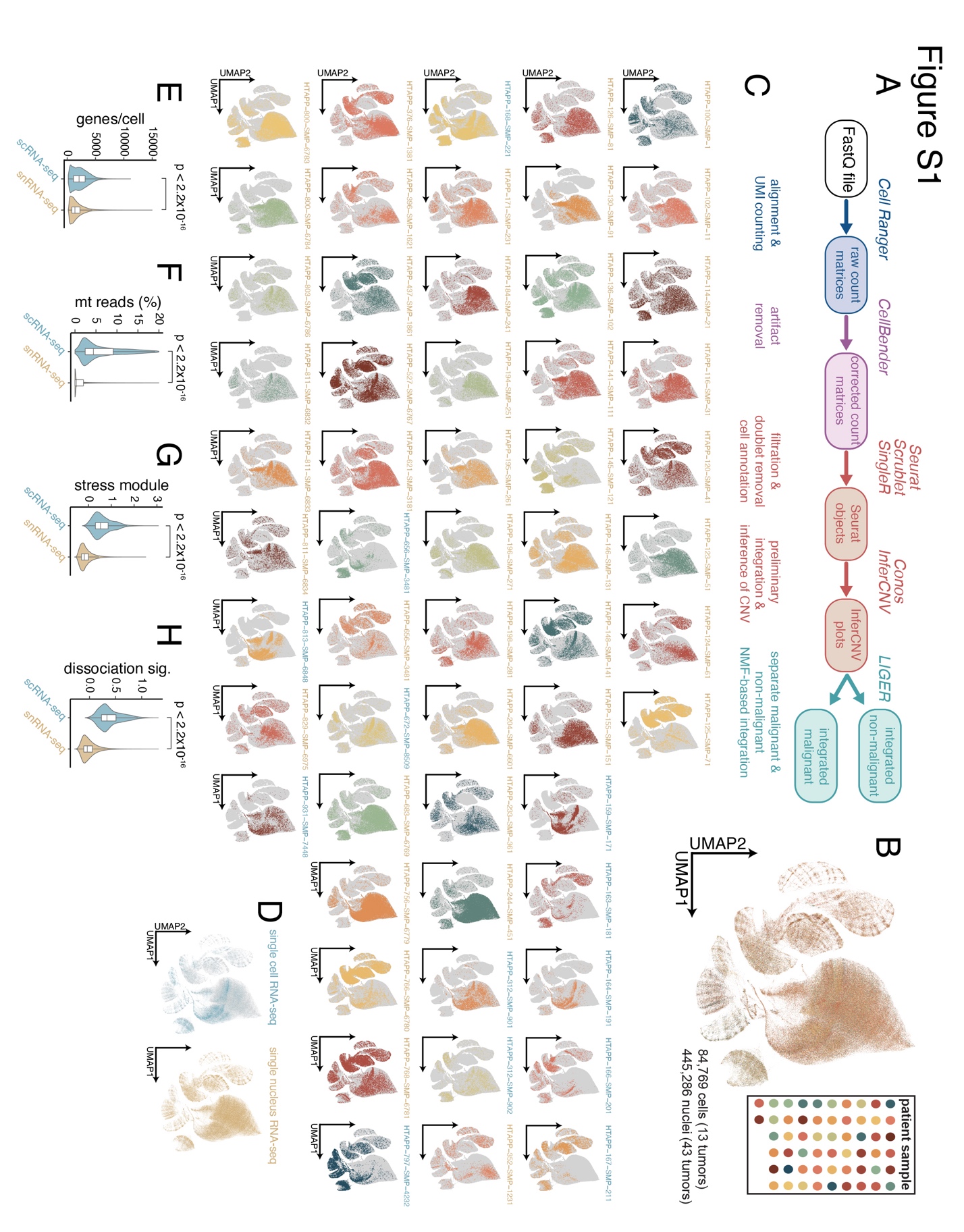

**Figure S2. Malignant cell type identification and copy-number inference, related to Figure 2.**

(A-C) UMAP plots of the neuroblastoma transcriptomic atlas, colored based on expression of *PTPRC* (A), *PECAM1* (B), and *NCAM1* (C).

(D) UMAP plot of one neuroblastoma sample, HTAPP-124-SMP-61, generated using snRNA-seq. Nuclei are colored based on annotated cell type. Malignant cell clusters are highlighted with grey shading.

(E) Inference of copy number variation for HTAPP-124-SMP-61. This heatmap is oriented by chromosomal position (columns) and individual cell (rows). Amplified regions are depicted with shades of red, while deletions are depicted in shades of blue. Immune and endothelial cells were used as reference cells (top).

(F) Bar plots showing cell type composition of the HTAPP neuroblastoma atlas, oriented to have all scRNA-seq datasets on the left and all snRNA-seq datasets on the right. Sample HTAPP-656-SMP-7481, which underwent both processing methods, is shown in the center.

(G) Micrographs of one HTAPP sample, HTAPP-171-SMP-231, stained by hematoxylin & eosin (H&E), and 3 immunohistochemical markers: PHOX2B, vimentin (VIM) and Ki67.

(H,I) Scatter plots comparing the proportion of cells in the SYMP or ADRN cell state (x-axis) to immunohistochemical stains for Ki-67 or PHOX2B (y-axis).

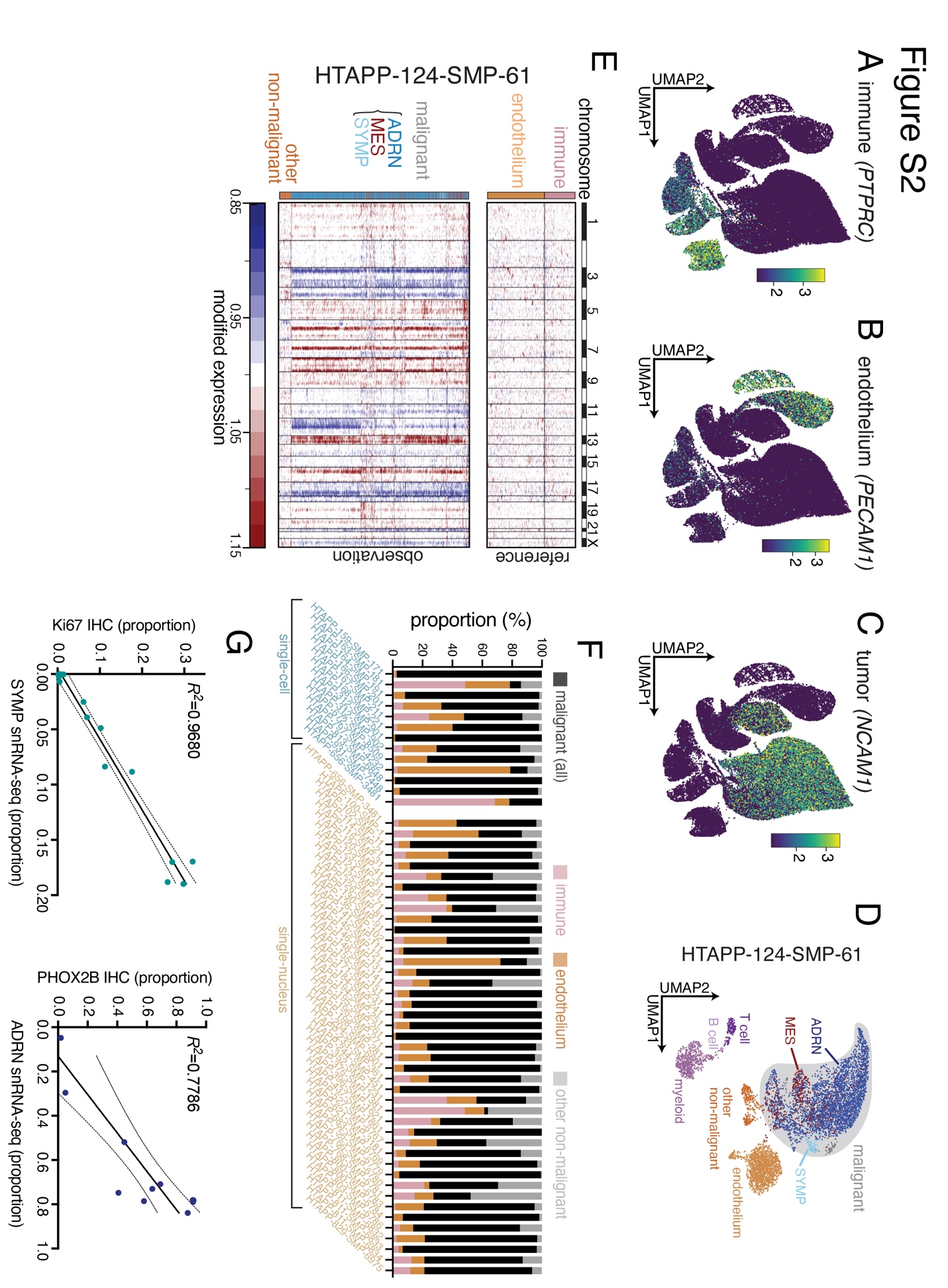

**Figure S3. Gene expression associated with malignant cell states and relationship to the fetal adrenal medulla, related to Figure 3.**

(A) UMAP plot of a reference fetal adrenal medulla from Jansky, et al.^1^ with annotated developmental states.

(B) Similarity scores of malignant cells/nuclei within the HTAPP neuroblastoma cohort when correlated to the reference adrenal medulla in (A).

(C) Comparison of late SCP similarity score from (C) to MES (left) and ADRN scores reported in van Groningen, et al.^2^

(D) UMAP plot of an alternate reference fetal adrenal medulla from Kildisiute, et al.^3^ with annotated developmental states.

(E) UMAP plot of HTAPP malignant cells/nuclei, colored based on predicted similarity to fetal adrenal medulla cell states from (D).

(F) Bar plots showing cell type distribution across samples within the HTAPP cohort, including non-malignant cell types (*top*) or subset to focus only on the malignant cell types in the dataset (*bottom*).

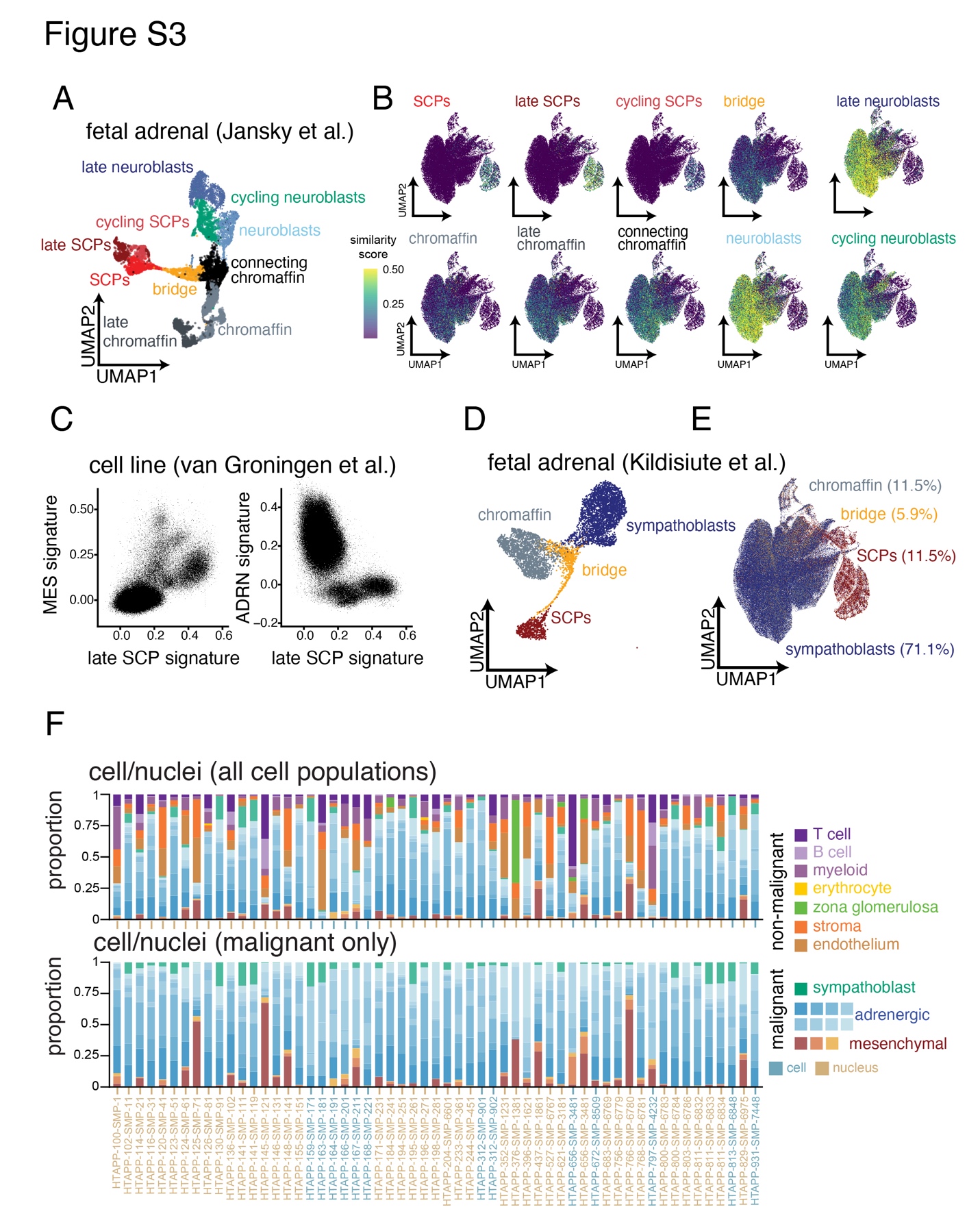

**Figure S4. Comparison of malignant cell states across clinical and molecular covariates, related to Figure 4.**

(A-D) Box plot of the proportion of malignant cells/nuclei within each developmental state, divided by MYCN status (A), Children’s Oncology Group (COG) risk stratum (B), age (C), and treatment status (D). For panel (D), we restricted analysis to intermediate and high-risk samples. Statistically credible differences, as determined by Bayesian composition analysis^4^, are shown with asterisks. Data are presented as median ± interquartile range.

(E) Bar plots showing the distribution of cell types across the HTAPP cohort, including non-malignant cell types (*top*) or subset to focus only on the malignant cell types in the dataset (*bottom*). Samples are organized by methylation grouping.

(F) Bar plots showing the distribution of cell types across the NCI TARGET neuroblastoma dataset, as estimated by CIBERSORTx bulk deconvolution. All cell types, including non-malignant cells are shown on top, and a comparison of only malignant cell types are shown on the bottom. Samples are organized by methylation grouping.

(G) Table summarizing clinical, developmental state composition, and survival of the 4 neuroblastoma methylation groups.

(H) Volcano plot of signaling pathway inference comparing group II adrenergic cells to adrenergic cells from the other methylation groupings.

(I) HALLMARK pathway analysis of statistically significant results from (H). No pathways were significantly enriched in group II adrenergic cells.

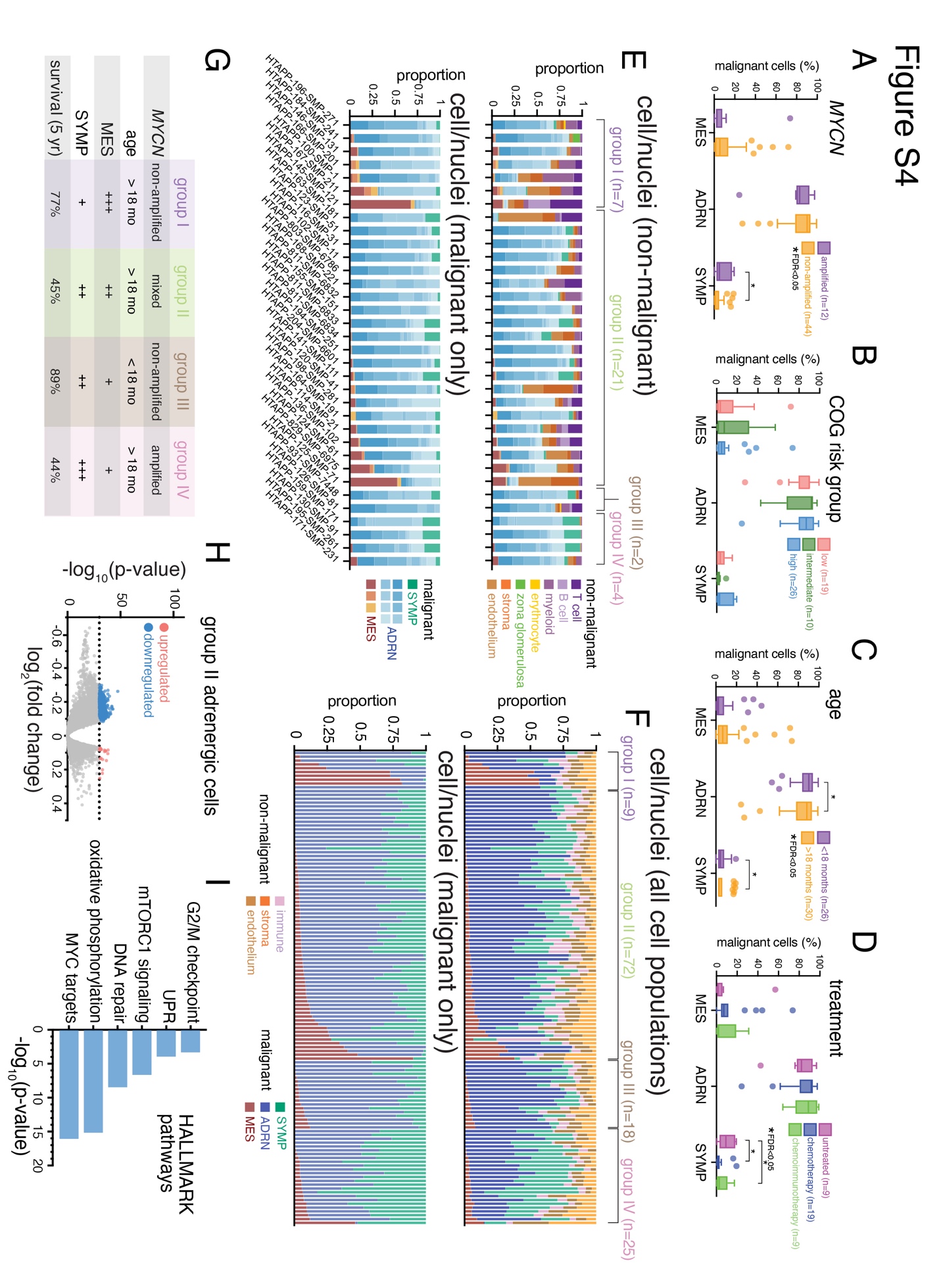

**Figure S5. Variation of immune cell type diversity across neuroblastoma, related to Figure 5.**

(A) UMAP plot of all immune cell/nuclei data colored based on dissociation method (fresh single-cell versus frozen single-nucleus, *top*) or by sample (*bottom*).

(B) Quality control metrics comparing immune scRNA-seq and snRNA-seq data. p-values calculated using the Wilcoxon signed rank test.

(C) UMAP plot of all immune cell/nuclei nuclei from the HTAPP neuroblastoma dataset, colored by patient sample.

(D,E) UMAP plots of T and NK nuclei from the HTAPP neuroblastoma dataset. Nuclei are colored based on the expression of cell markers (D) or originating patient sample (E).

(F) Dot plot showing expression of T/NK markers in each annotated cluster. The radius of each dot reflects the percentage of nuclei for which marker expression was detected, and the color of each circle denotes the expression level.

(G) Bar plots showing cell type proportions of T/NK subtypes within the HTAPP cohort.

(H,I) UMAP plots of B cell and plasma cell snRNA-seq data. Nuclei are colored based on the expression of cell markers (H) or originating patient sample (I).

(J) Dot plot showing expression of B cell markers in each annotated cluster. The radius of each dot reflects the percentage of nuclei for which marker expression was detected, and the color of each circle denotes the expression level.

(K) Bar plots showing cell type proportions of B cell subtypes within the HTAPP cohort.

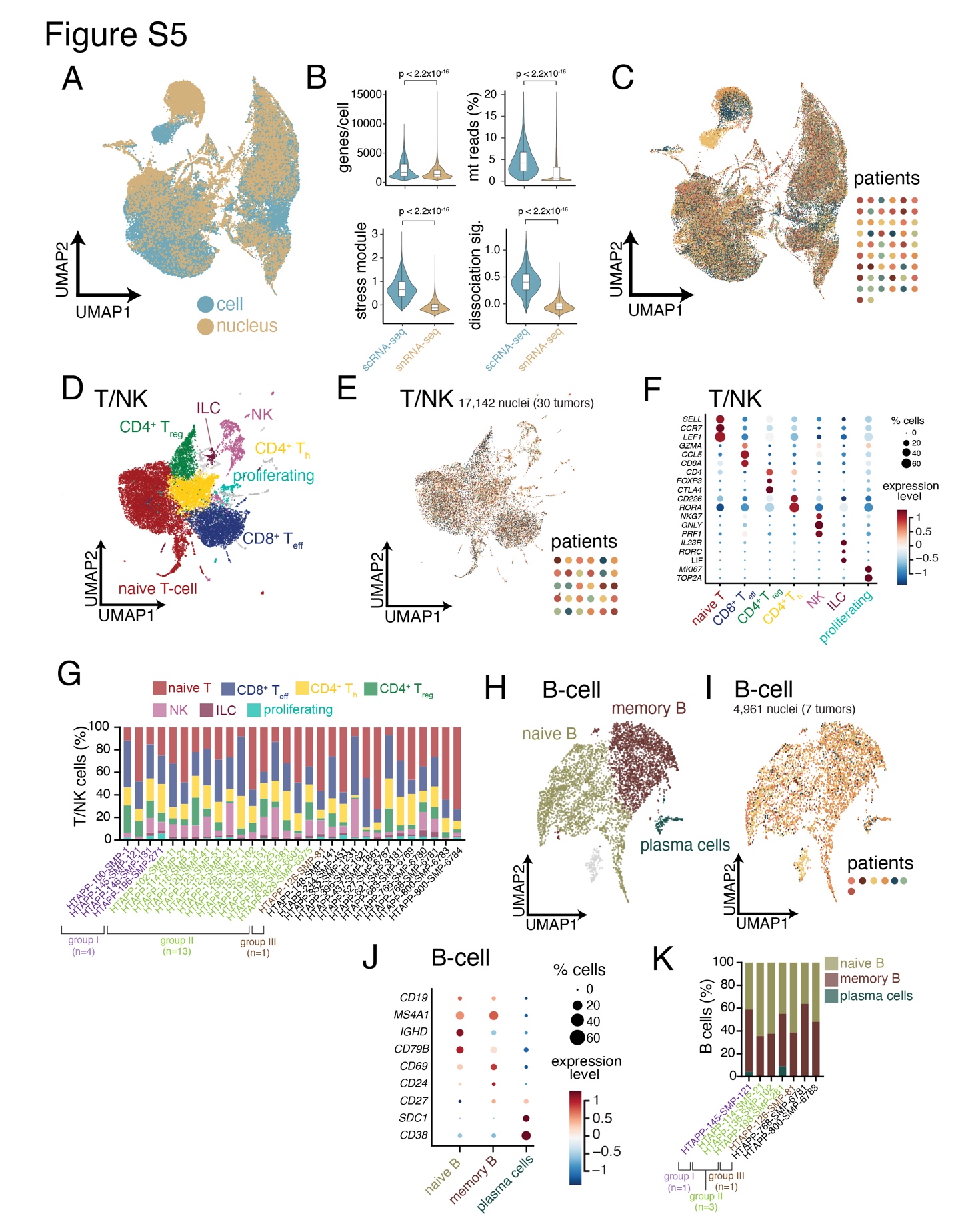

**Figure S6. CODEX validation and data exploration, related to Figure 7.**

(A) Heatmap of annotated cell types (rows), based on detection of markers (columns). Multiple immune cell types were readily identified. Tumor and stroma were defined as cells without immune cell marker positivity.

(B) Heatmap showing subdivision and further annotation of the tumor and stroma cell type from (A). These annotations were based primarily using CD56 (NCAM1), collagen IV (COL4), HLA-DR, podoplanin and vimentin.

(C) Montage of a single region from a portion of one CODEX dataset. Each CODEX sample was imaged using a cocktail of 52 oligonucleotide-conjugated antibodies (Table S8). Each antibody is depicted in false grey color.

(D-E) Sequential sections of a CODEX sample showing H&E staining (D) and a 4-color overlay of tumor and stromal markers (E) showing CD56 (red), podoplanin (green), collagen IV (white) and vimentin (blue).

(F) Prediction of cell identities using co-registered H&E and CODEX images. In this panel, 4 cell types (CD56+VIM+COL4+PDPN–, CD56+VIM+COL4–PDPN–, VIM+COL4+ stroma, and classical dendritic cells).

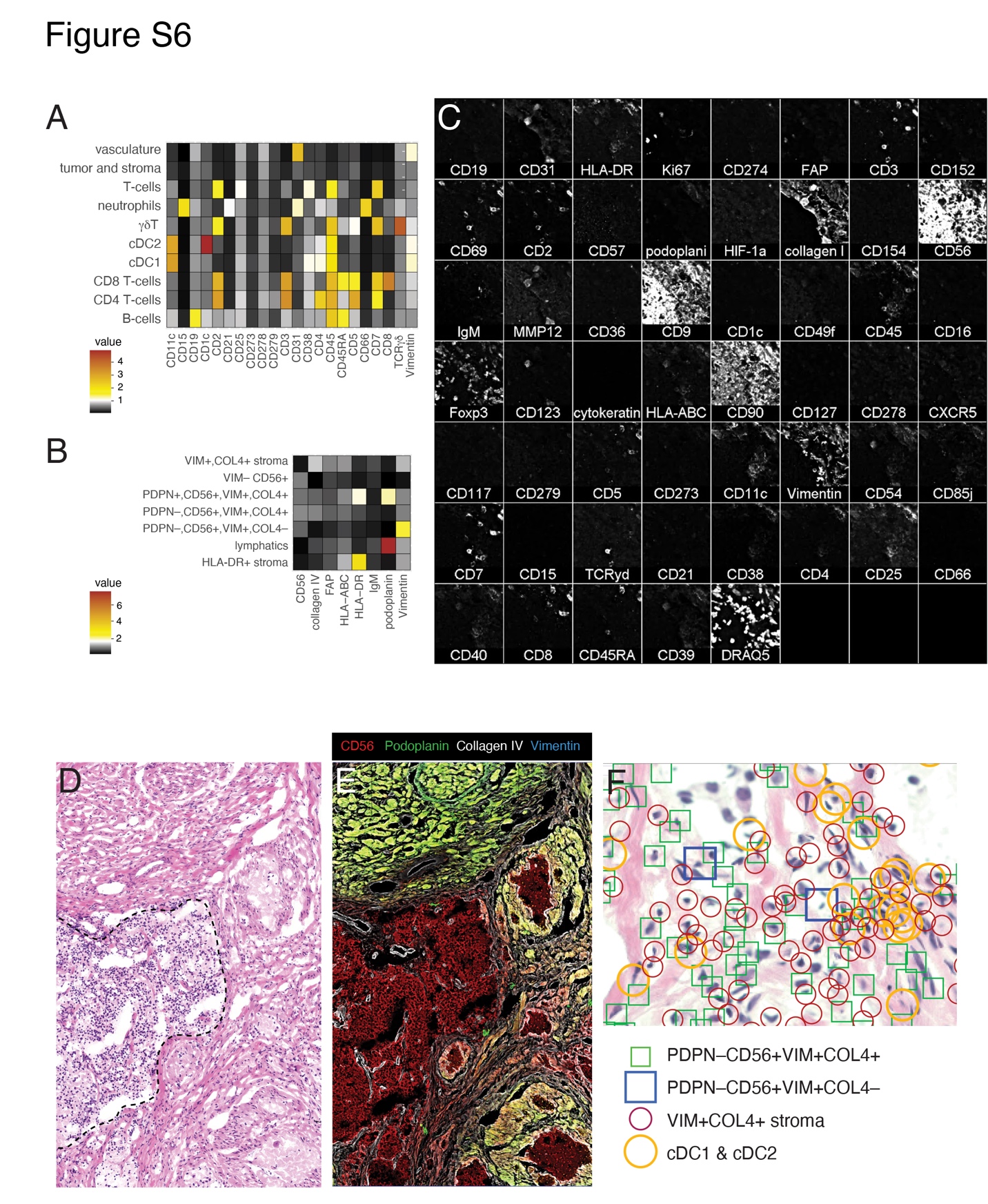

**METHODS**

**KEY RESOURCES TABLE**

| **Reagent or Resource** | **Source** | **Identifier** |
| --- | --- | --- |
| *Antibodies* | | |
| anti-PHOX2B | Abcam | ab183741; RRID:AB_2857845 |
| anti-Vimentin | Roche | 790-2917; RRID:AB_2687607 |
| anti-Ki67 | Thermo Fischer | RM-9106;  RRID: AB_2341197 |
| anti-CD3 | Biocare Medical | CME324B;  RRID:AB_2923282 |
| anti-CD8 | Roche | 790-4460;  RRID:AB_2335985 |
| anti-CD68 | Roche | 790-2931;  RRID:AB_2335972 |
| anti-CD163 | Roche | 760-4437;  RRID:AB_2335969 |
| Metal-conjugated antibodies for MIBI | various sources | see Table S8 |
| Oligonucleotide-conjugated antibodies for CODEX | various sources | see Table S8 |
| *Biological samples* | | |
| Patient neuroblastoma tissue | St. Jude Children’s Research Hospital | see Table S1 |
| Patient neuroblastoma tissue | Dana-Farber Cancer Institute | see Table S1 |
| Flash-frozen orthotopic patient-derived xenograft tissue | St. Jude Children’s Research Hospital Childhood Solid Tumor Network | <https://cstn.stjude.cloud/> |
| *Chemicals, Peptides and Recombinant Proteins* | | |
| Papain Dissociation Kit | Worthington Biochemical Corporation | LK003150 |
| Maxima H Minus reverse transcriptase and 5x Maxima RT buffer | Thermo Fisher Scientific | EP0751 |
| NxGen RNase inhibitor | Lucigen | 30821 |
| 10 mM dNTP solution | New England Biolabs | N0477L |
| Proteinase K | New England Biolabs | P81075 |
| Exonuclease I (E. coli) | New England Biolabs | M0293 |
| Klenow enzyme | New England Biolabs | M0210 |
| Terra Direct PCR mix | Takara Biosciences | 639270 |
| AMPure XP beads | Beckman Coulter | A63880 |
| EB solution | Qiagen | 19086 |
| Nextera XT kit | Illumina | FC-131-1096 |
| *Critical Commercial Assays* | | |
| Chromium Single Cell 3’ Library & Gel Bead Kit v2 | 10x Genomics | 120237 |
| Chromium Single Cell 3’ Library & Gel Bead Kit v3 | 10x Genomics | 1000075 |
| Chromium Single Cell A Chip kit | 10x Genomics | 120236 |
| Chromium Single Cell B Chip kit | 10x Genomics | 1000153 |
| Chromium i7 Multiplex Kit | 10x Genomics | 120262 |
| DISCOVERY Rhodamine 6G detection kit | Roche Tissue Diagnostics | 760-244 |
| DISCOVERY DCC detection kit (coumarin) | Roche Tissue Diagnostics | 760-240 |
| DISCOVERY Red 610 detection kit | Roche Tissue Diagnostics | 760-245 |
| DISCOVERY FAM detection kit | Roche Tissue Diagnostics | 760-243 |
| DISCOVERY Cy5 detection kit | Roche Tissue Diagnostics | 760-238 |
| DISCOVERY OmniMap anti-Rb HRP | Roche Tissue Diagnostics | 760-4311 |
| DISCOVERY OmniMap anti-Ms HRP | Roche Tissue Diagnostics | 760-150 |
| Maxwell RSC DNA FFPE kit | Promega | AS1450 |
| Infinium MethylationEPIC BeadChip Kit | Illumina | 20042130 |
| Quant-iT PicoGreen dsDNA assay | Thermo Fisher Scientific | P7589 |
| TruSeq DNA Exome Library Prep Kit | Illumina | 20020615 |
| TruSeq DNA Rapid Exome Library Prep Kit | Illumina | 20020616 |
| Maxpar X8 Multimetal Labeling Kit | Fluidigm | 201300 |
| *Recombinant DNA* | | |
| Slide-seqV2 template switch oligo | IDT | 5’-AAGCAGTGGTATCAACGCAGAGTGAATrG+GrG-3’ |
| dN-SMRT oligonucleotide | IDT | 5’-AAGCAGTGGTATCAACGCAGAGTGANNNGGNNNB-3’ |
| TruSeq PCR handle primer | IDT | 5’-CTACACGACGCTCTTCCGATCT-3’ |
| SMART PCR primer | IDT | 5’-AAGCAGTGGTATCAACGCAGAGT-3’ |
| TruSeq5 sequencing primer | IDT | 5’-AATGATACGGCGACCACCGAGATCTACACTCTTTCCCTACACGACGCTCTT  CCGATCT-3’ |
| *Deposited Data* | | |
| Single-cell dissociation protocol | Slyper et al., 2020^5^ | <https://www.protocols.io/view/htapp-dissociation-of-human-neuroblastoma-tumors-t-98ah9se> |
| Single-nucleus dissociation protocol (TST) | Slyper et al., 2020^5^ | <https://www.protocols.io/view/htapp-tst-nuclei-isolation-from-frozen-tissue-bhbcj2iw> |
| Whole exome sequencing methods (St. Jude Children’s Research Hospital protocol) | This paper | <https://www.protocols.io/view/exome-whole-genome-and-bulk-rna-sequencing-st-jude-bvvfn63n> |
| Whole exome sequencing methods (Broad Institute protocol) | This paper | <https://www.protocols.io/view/exome-ice-sequencing-methods-the-broad-institute-bxwvppe6> |
| Single-cell/nucleus RNA-sequencing data | This paper | <https://humantumoratlas.org/>; <https://viz.stjude.cloud/community/human-tumor-atlas-network-consortium~6> |
| Whole exome sequencing data | This paper | <https://humantumoratlas.org/> |
| Bulk RNA-sequencing data | This paper | <https://humantumoratlas.org/> |
| Illumina MethylationEPIC 850k data | This paper | GEO GSE216650 |
| MIBI imaging data | This paper | <https://humantumoratlas.org/> |
| CODEX imaging data | This paper | <https://humantumoratlas.org/> |
| Slide-SeqV2 imaging data | This paper | <https://humantumoratlas.org/> |
| TARGET methylation data | TARGET project | <https://ocg.cancer.gov/programs/target> |
| TARGET bulk RNA-seq data | TARGET project | <https://ocg.cancer.gov/programs/target> |
| Human fetal adrenal gland single-cell RNA-seq data | Jansky et al., 2021^1^ | <https://adrenal.kitz-heidelberg.de/developmental_programs_NB_viz/> |
| Human fetal adrenal gland single-cell RNA-seq data | Kildisiute et al., 2021^3^ | <https://www.neuroblastomacellatlas.org/> |
| *Software and Algorithms* | | |
| Cell Ranger v3.0 | 10x Genomics | <https://support.10xgenomics.com/single-cell-gene-expression/software/downloads/latest> |
| Cellbender v0.2.0 | Fleming et al., 2022^6^ | <https://github.com/broadinstitute/CellBender> |
| Scrublet | Wolock et al., 2019^7^ | <https://github.com/AllonKleinLab/scrublet> |
| Python | Python Software Foundation | <https://www.python.org/> |
| R | The R Foundation | <https://www.r-project.org/> |
| RStudio | RStudio | <https://rstudio.com/> |
| InferCNV | Trinity CTAT Project | <https://github.com/broadinstitute/inferCNV/wiki> |
| BioCircos | Cui et al., 2016^8^ | <https://github.com/cran/BioCircos> |
| Seurat | Hao et al., 2021^9^ | <https://satijalab.org/seurat/> |
| Conos | Barkas et al., 2019^10^ | <https://github.com/kharchenkolab/conos> |
| LIGER | Welch et al., 2019^11^ | <https://github.com/MacoskoLab/liger> |
| SingleR | Aran et al., 2019^12^ | <https://bioconductor.org/packages/release/bioc/html/SingleR.html> |
| ComplexHeatmap | Gu et al., 2016^13^ | <https://bioconductor.org/packages/release/bioc/html/ComplexHeatmap.html> |
| pySCENIC v0.11.2 | Van de Sande et al., 2020^14^ | <https://scenic.aertslab.org/> |
| scCODA v0.1.2post1 | Büttner et al., 2021^4^ | <https://github.com/theislab/scCODA> |
| CellTypist v1.0.0 | Domínguez Conde et al., 2022 | <https://www.celltypist.org/> |
| SCpubr v0.1.0 | Blanco-Carmona, 2022^15^ | <https://github.com/enblacar/SCpubr> |
| Rtsne v0.15 | Van der Maaten and Hinton, 2008^16^ | <https://github.com/jkrijthe/Rtsne> |
| ConsensusClusterPlus v1.54.0 | Wilkerson and Hayes, 2010^17^ | <https://bioconductor.org/packages/ConsensusClusterPlus> |
| DMRcate v1.18.0 | Peters et al., 2015^18^ | <https://bioconductor.org/packages/release/bioc/html/DMRcate.html> |
| HALO v3.2.1851.354 | Indica Labs | <https://indicalab.com/halo/> |
| BWA-aln | Li and Durbin, 2009^19^ | <http://bio-bwa.sourceforge.net/> |
| Bambino | Edmonson et al., 2011^20^ | <https://github.com/NCIP/cgr-bambino> |
| CrossMap v0.2.4 | Zhao et al., 2014^21^ | <https://github.com/liguowang/CrossMap> |
| Mutect2 v4.1.2.0 | Cibulskis et al., 2013^22^ | <https://gatk.broadinstitute.org/hc/en-us/articles/360037593851-Mutect2> |
| SomaticSniper v1.0.5.0 | Larson et al., 2012^23^ | <http://gmt.genome.wustl.edu/packages/somatic-sniper/> |
| VarScan2 v2.4.3 | Koboldt et al., 2012^24^ | <http://varscan.sourceforge.net/> |
| MuSE v1.0rc | Fan et al., 2016^25^ | <https://github.com/danielfan/MuSE> |
| Strelka2 v2.9.10 | Kim et al., 2018^26^ | <https://github.com/Illumina/strelka> |
| Annovar | Wang et al., 2010^27^ | <https://annovar.openbioinformatics.org/en/latest/> |
| NetBID | Du et al., 2018^28^ | <https://github.com/jyyulab/NetBID> |
| SJARACNe | Khatamian et al., 2019^29^ | <https://github.com/jyyulab/SJARACNe> |
| CIBERSORTx | Newman et al., 2019^30^ | <https://cibersortx.stanford.edu/> |
| Enrichr | Kuleshov et al., 2016^31^ | <https://maayanlab.cloud/Enrichr/> |
| bioformats2raw |  | <https://github.com/glencoesoftware/bioformats2raw> |
| raw2ometiff |  | <https://github.com/glencoesoftware/raw2ometiff> |
| ilastik | Berg et al., 2019^32^ | <https://www.ilastik.org/> |
| MIBI Analysis tools | Keren et al., 2018^33^ | <https://github.com/lkeren/MIBIAnalysis> |
| MIBIStitch |  | <https://github.com/dtebaykin/MibiStitch> |
| deepcell-tf v0.6.0 | Greenwald et al., 2022; Van Valen et al., 2016^34,35^ | <https://github.com/vanvalenlab/deepcell-tf> |
| FlowSOM | Van Gassen et al., 2015^36^ | <https://bioconductor.org/packages/release/bioc/html/FlowSOM.html> |
| Marker enrichment modeling (MEM) | Diggins et al., 2017^37^ |  |
| QuPath v0.3.2 | Bankhead et al., 2017^38^ | <https://qupath.github.io/> |
| Microvolution |  | <http://www.microvolution.com/> |
| CODEX data analysis tools | Goltsev et al., 2018^39^ | <https://github.com/nolanlab/CODEX> |
| slideseq-tools | Stickels et al., 2021^40^ | <https://github.com/MacoskoLab/slideseq-tools> |
| RCTD | Cable et al., 2022^41^ | <https://github.com/dmcable/spacexr> |
| TANGRAM | Biancalani et al., 2021^42^ | <https://github.com/broadinstitute/Tangram> |
| Squidpy | Palla et al., 2022^43^ | <https://squidpy.readthedocs.io/en/stable/index.html> |
| CSIDE | Cable et al., 2022^44,45^ | <https://github.com/dmcable/spacexr> |
| GraphPad Prism v9 | GraphPad Software Inc. | <https://www.graphpad.com/scientific-software/prism/> |
| BioRender | BioRender | <https://biorender.com/> |

**RESOURCE AVAILABILITY**

Further information and requests for resources and reagents should be directed to and will be fulfilled by the lead contact, Dr. Michael Dyer.

**MATERIALS AVAILABILITY**

This study did not generate new unique reagents.

**EXPERIMENTAL MODELS AND SUBJECT DETAILS**

**Human subjects**

The accrual of patient tumor material for this study underwent IRB approval at the institutional site of origin. St. Jude Children’s Research Hospital IRB approved protocols XPD09-234 and XPD18-051. Dana-Farber Cancer Institute IRB approved protocols 11-104 and 17-104.

**Orthotopic patient-derived xenograft (O-PDX) tissue**

Frozen tissue from orthotopic patient-derived xenografts (O-PDXs) were obtained through the St. Jude Childhood Solid Tumor Network (<https://cstn.stjude.cloud/>)^46^.

**METHOD DETAIL**

**Sample processing for single-cell RNA-seq**

Fresh tissue samples were immediately processed using the Papain Dissociation System (Worthington), which we had previously identified as the optimal dissociation method for neuroblastoma^5^. Briefly, small fragments (<500 mg) were rinsed with phosphate-buffered saline without calcium or magnesium (PBS), minced using a scalpel, and then incubated in 5 mL papain-DNase I solution at 37°C for 15-30 min. The dissociated tumors were triturated, filtered through a 40 μm strainer, and centrifuged at 500 x *g* for 5 min. The cell pellets were then resuspended in 3.1 mL resuspension buffer (2.7 mL Earle’s buffered saline solution, 300 μL albumin-ovomucoid inhibitor, 150 μL DNase I). To remove debris, the cell suspensions were then carefully layered over 5 mL albumin-ovomucoid inhibitor cushion and centrifuged at 100x*g* for 6 min. The resulting pellet was resuspended in PBS and filtered through a 40 μm strainer to generate single-cell suspensions for scRNA-seq. Cell viability was confirmed using trypan blue dye exclusion.

**Sample processing for single-nucleus RNA-sequencing**

To generate flash-frozen tissue, resected fresh tumor tissue fragments were immediately placed on dry ice and stored at -80°C. Generation of single-nucleus suspensions from flash-frozen tissue using the TST protocol were described previously^5^. At the time of processing, frozen fragments were manually dissociated using Noyes spring scissors in 1 mL TST buffer (73 mM sodium chloride, 5 mM Tris [pH 8.0], 0.5 mM calcium chloride, 10.5 mM magnesium chloride, 0.01% bovine serum albumin, 0.03% Tween-20). Nuclear suspensions were filtered through a 40 μm strainer and diluted with an additional 1 mL TST buffer and 3 mL ST buffer (73 mM sodium chloride, 5 mM Tris [pH 8.0], 0.5 mM calcium chloride, 10.5 mM magnesium chloride). The cell suspension was centrifuged at 500x*g* at 4°C for 5 min. The nuclear pellet was resuspended in ST buffer and filtered through a 40 μm strainer to generate single-nuclear suspensions for snRNA-seq. Appropriate nucleus extraction was confirmed via manual inspection of Trypan blue stained nuclei suspension.

**Generation of single-cell/nucleus RNA-seq libraries**

Single cell or nuclear suspensions were performed using 10x Genomics Single-cell 3’-RNA Expression Solution kits version 2 or 3. Ten-thousand cells or nuclei were input into the Chromium controller (10x Genomics) for microfluidic partitioning followed by barcoded reverse transcription. Subsequent cDNA amplification, fragmentation, adapter ligation and sample indexing were performed according to manufacturer instructions. Libraries were pooled and sequenced on either an Illumina NextSeq or Illumina NovaSeq 6000 sequencer with paired-end reads as follows: read 1 - 26 nucleotides, read 2 - 55 nucleotides, index 1 - 8 nucleotides, index 2 - 0 nucleotides.

**Data analysis of single-cell/nucleus RNA-seq data**

We pre-processed and performed initial analyses of single-cell and single-nucleus RNA-seq (sc/snRNA-seq) data using our previously developed toolbox for fresh and frozen human tumors, implemented within Cumulus^5^ in the Terra Cloud platform (<https://app.terra.bio/>). We demultiplexed raw sequencing reads into FASTQ files using Cell Ranger mkfastq v3.0 (10x Genomics) and we aligned reads and quantified gene counts as UMIs using Cell Ranger count v3.0 (10x Genomics). For scRNA-seq, we aligned reads to the human GRCh38 genome. For snRNA-seq, we aligned reads to a ‘pre-mRNA’ human GRCh38 reference, which included reads that mapped to introns^5^.

Gene and cell quality control consisted of three steps: (i.) removal of noise due to ambient RNA, (ii.) removal of cell doublets, and (iii.) removal of cells with low library complexity or number of genes detected. We used CellBender remove-background to remove background noise counts from the gene count matrices due to ambient RNA.^6^ For each dataset, we adjusted the CellBender parameters “expected-cells” and “total-droplets-included” based on the barcode rank plot. We kept all other settings, including the number of epochs and learning rate, at their default values, and used a false positive rate of 0.01 when controlling for signal to noise counts. We visually inspected the ELBO plot to verify the inference procedure had converged. In downstream analyses, we used the final count matrix file (FPR_0.01_filtered.h5) with ambient RNA counts subtracted and only droplets with a > 50% posterior probability of containing cells.

For each individual sample’s gene count matrix, we used Scrublet to identify potential cell or nucleus doublets^7^. We calculated doublet scores for each cell and assigned cells as potential doublets if they exceeded the automatically calculated threshold score. After data integration and clustering all cells across all samples (described in more detail below), we inspected each cell cluster. Cell clusters where the majority of cells in the cluster had been assigned as potential doublets and the differentially expressed genes of the cluster were consistent with multiple cell types were removed from subsequent analyses. We performed this doublet filtering separately for the scRNA-seq and snRNA-seq data modalities. Overall, fewer than 5% of all cells or nuclei were removed as potential doublets.

We removed low-quality cells and nuclei with a low number of UMIs or genes detected. Depending on the data modality (single cell or single nucleus) and the 10x chemistry (V2 or V3), we used different thresholds. For single nucleus data collected using V2 chemistry, we kept nuclei with at least 400 UMIs and 200 genes detected. For all other data, we kept cells or nuclei with at least 1000 UMIs and 500 genes detected. Finally, we filtered out those cells with greater than 20% of UMIs mapped to mitochondrial genes. In addition to the above quality controls, we also removed nine patient samples (from an initial cohort of 64 samples) for the following reasons: poor overall quality control statistics (<10% of reads mapped to the genome or fewer than 400 cells), disagreements across technical replicates, and absence of distinct cell subsets.

We carried out data integration of all transcriptomic data to assign cells to broad lineages: immune, endothelial, stromal/malignant, parenchyma, or technical doublet. We performed integration for scRNA-seq and snRNA-seq modalities separately but combined the modalities in subsequent analyses. To integrate the patient samples, we ran Conos^10^ with default settings, building the joint graph in principal component analysis (PCA) space. We then performed initial annotation of the cell lineages using a combination of differentially expressed genes, known neuroblastoma gene signatures, and the automated annotation package SingleR, as previously described^5,12^.

We clustered cells in each individual tumor sample to separate malignant cells from non-malignant cells, which was required for our later analysis of malignant cell states. For each sample, we carried out data normalization, feature selection, dimensionality reduction, clustering, and visualization using our previously described Cumulus workflow on Terra^5^. To identify the malignant cells within each sample, we inferred chromosomal copy-number aberrations (CNAs) from the gene-expression data using inferCNV as previously described^5^. We used immune cells and endothelial cells as a healthy cell reference to infer CNAs in the remaining cells, which were a mix of malignant cells and stromal cells. As neuroblastoma cells share several transcriptional programs with stromal cells, the presence of CNAs more clearly distinguishes these cell subsets. We used both the presence of CNAs and expression of neural crest differentiation programs in assigning malignant cells. BioCircos plots from samples (Figure S2F) were generated using custom code.

To identify cell types and states within the immune and malignant cell lineages, we applied the non-negative matrix factorization algorithm LIGER^11^, which allows simultaneous data integration across patient samples and discovery of shared cell states across samples. We combined samples from both scRNA-seq and snRNA-seq modalities, and for both the immune and malignant cell lineages, we used twenty factors in the factorization. To interpret the different cell types and states represented by each factor, we examined the top 100 genes loading each factor. We further subdivided the immune cells into T/NK, B, and myeloid cell lineages based on the factors and based on predicted annotations from CellTypist using the Immune_All_Low and Immune_All_High models^47^. We then sub-clustered the T/NK, B, and myeloid cell lineages using LIGER with eight, four, and four factors respectively.

Processing of single-nucleus RNA-seq data from orthotopic patient-derived xenografts (O-PDXs) were generated in an identical manner to data from human tumor tissue, with the exception that sequenced reads were aligned to a combined human and mouse reference (GRCh38 and mm10, respectively; available at <https://support.10xgenomics.com/single-cell-gene-expression/software/downloads/latest>). Nuclei that mapped to the mouse genome were removed from downstream analysis. For comparison of the patient tumor to O-PDXs, we used Seurat’s anchor-based transfer method^9^. Specifically, we used the MapQuery workflow within Seurat, using anchors defined by the reciprocal PCA reduction method.

Stress and dissociation scores (Fig. S1E) were generated using Seurat’s AddModuleScore. The stress signature was obtained from Table S1 of Biemann, et al.^48^, and the dissociation signature was obtained from Supplementary Table 5 of van der Brink, et al.^49^

To compare malignant neuroblastoma sc/snRNA-seq data, we downloaded scRNA-seq data from human the fetal adrenal medulla from Jansky, et al^1^ (available at: <https://adrenal.kitz-heidelberg.de/developmental_programs_NB_viz/>) or Kildisiute, et al.^3^ (available at: <https://neuroblastoma-cell-atlas.cog.sanger.ac.uk/adr_all.rds>). The SingleR package was used to generate assignment scores based on the Spearmann correlation between experimental data (HTAPP neuroblastoma) and reference annotations (from either fetal adrenal dataset)^12^.

For estimation of transcription factor, we applied pySCENIC to scRNA-seq data from malignant neuroblastoma cells using default parameters^14,50^. First, we generated inferred gene regulatory networks using the ‘pyscenic grn’ command using a list of curated transcription factors (TFs; available at: <https://github.com/aertslab/pySCENIC/blob/master/resources/hs_hgnc_tfs.txt>). We then used the ‘pyscenic ctx’ command without the dropout masking option to refine regulons for those genes that do not have an adjective TF binding motifs (based on previously defined cis-regulatory footprinting, available at: <https://resources.aertslab.org/cistarget/motif2tf/motifs-v10nr_clust-nr.hgnc-m0.001-o0.0.tbl>). Finally, we used the ‘pyscenic aucell’ command to score individual cells for inferred TF activity. Results from pySCENIC were then imported and combined with the scRNA-seq expression matrix within Seurat. Wilcoxon rank sum testing was performed to identify those TFs that were predicted to have differential activity between the 3 malignant cell types.

**Bulk sequencing for WES and somatic variant calling**

Complete details for methods used for whole exome data generation are available for the protocols used at St. Jude Children’s Research Hospital and Broad Institute, at [dx.doi.org/10.17504/protocols.io.bvvfn63n](https://dx.doi.org/10.17504/protocols.io.bvvfn63n) and [dx.doi.org/10.17504/protocols.io.bxwvppe6](https://dx.doi.org/10.17504/protocols.io.bxwvppe6). Genomic DNA was acoustically sheared using a Covaris LE220 ultrasonicator. Library preparation and exome target enrichment was performed using the TruSeq DNA Exome Library Prep Kit (St. Jude Children’s Research Hospital) or the TruSeq DNA Rapid Exome Library Prep Kit (Broad Institute), according to the manufacturer’s instructions. Paired-end 100 cycle sequencing was performed on a NovaSeq 6000 (St. Jude Children’s Research Hospital) or Illumina HiSeq 4000 (Broad Institute) sequencer.

Whole-exome sequencing (WES) reads of samples from St. Jude Children’s Research Hospital were mapped to GRCh37-lite by BWA-aln. Somatic SNVs and INDELs were called on matched tumor/normal samples by Bambino as previously described^51,52^. The coordinates of the somatic SNVs and INDELs were then lifted over to GRCh38 by CrossMap^21^.

WES reads of the samples from Dana-Farber Cancer Institute were mapped to GRCh38 by BWA-mem. Somatic SNVs and INDELs were called on matched tumor/normal samples by five callers: Mutect2^22^, SomaticSniper^23^, VarScan2^24^, MuSE^25^, and Strelka2^26^, followed by consensus calling. Multiple filtering steps were then applied to exclude potential artifacts. We kept variants that fulfilled the following criteria: coverage depth of the variant > 10, variant allele frequency > 0.02, alternative allele count ≥ 4, allele population frequency in public databases < 0.01 (gnomadAD, 1000 genomes, ExAC and Exome Sequencing Projects), mappability > 0.7, and not co-localized with repeat elements and GC percentage between 0.4 and 0.6. Variant annotation was performed using Annovar^27^. For sequencing results obtained from formalin-fixed samples, an additional round of filtration was performed to remove potential FFPE artifacts from formaldehyde deamination of cytosines (C > T); to accomplish this, we ran Mutect2 tumor-only mode to identify those variants with orientation bias and removed them from the somatic variant list. The variants in Table S2 are reported either as an unfiltered list or as a further curated list that includes only protein altering mutations (including missense, nonsense, frameshift, in-frame, and splice mutations) from those NBL signature genes that were previously reported in Brady et al.^53^, or through the Pediatric Cancer Genome Project (available at: <https://pecan.stjude.cloud/proteinpaint/heatmap/ST,NBL>).

**Immunohistochemistry**

Tissues were fixed in 10% neutral buffered formalin, embedded in paraffin, sectioned at 4-μm, mounted on positive charged glass slides (Superfrost Plus; 12-550-15, Thermo Fisher Scientific, Waltham, MA) that were dried at 60°C for 20 minutes. The following immunohistochemistry protocols were used on the Ventana DISCOVERY ULTRA automated stainer (Ventana Medical Systems, Inc., Tucson, AZ):

1. PHOX2B staining: heat-induced epitope retrieval with cell conditioning media 1 (Ventana Medical Systems) for 64 minutes, then primary antibody was applied for 60 min at a 1:100 dilution.
2. Vimentin staining: heat-induced epitope retrieval with cell conditioning media 1 (Ventana Medical Systems) for 32 minutes, then primary antibody (ready-to-use formulation) was applied 20 min.
3. Ki67 staining: heat-induced epitope retrieval with cell conditioning media 1 (Ventana Medical Systems) for 32 minutes, then primary antibody was applied for 60 min at a 1:200 dilution.

After primary antibody incubation, visualization was performed with DISCOVERY OmniMap anti-Rb HRP (Ventana Medical Systems), DISCOVERY ChromoMap DAB kit (Ventana Medical Systems), Hematoxylin II (Ventana Medical Systems), and Bluing reagent (Ventana Medical Systems). For automated cell counting, we scanned slides using a 3DHistech slide scanner at 40X magnification. We converted the 3DHistech MRXS images to ome.tiff using bioformats2raw and raw2ometiff command line tools. We used ilastik^32^, an interactive machine-learning segmentation software to segment DAB-positive pixels (stained as brown to dark brown) and Hemoxylin-positive pixels (nuclei stained as dark blue) at 10X magnification which is sufficient to see individual nuclei. For nuclear stains (Ki67 and PhOX2B), we calculated the index by ratio of the area of DAB-positive pixels to the sum of DAB-positive pixels and Hemoxylin-positive pixels.

**Multiplexed immunofluorescence**

All formalin-fixed, paraffin-embedded (FFPE) tissues were sectioned at 4-μm and mounted on positively charged glass slides. All tissue sections used for immunofluorescence labeling underwent deparaffinization followed by heat-induced antigen retrieval using prediluted Cell Conditioning Solution 1 (ULTRA CC1, Ventana Medical Systems) for 4 minutes. Primary antibodies were serially applied using the U DISCOVERY 5-Plex IF procedure and the following reagents and kits (all from Roche): DISCOVERY inhibitor, prediluted Cell Conditioning Solution 2, ready to use DISCOVERY OmniMap anti-Rb HRP, and visualization of PHOX2B using a DISCOVERY Rhodamine 6G kit, CD3 using a DISCOVERY DCC kit, CD8 using a DISCOVERY Red 610 kit, CD68 using a DISCOVERY CY5 Kit and CD163 using a DISCOVERY FAM Kit. Antibody dilution, clone ID, and incubation times are provided in Table S8. Coverslips were hand-mounted on slides with Prolong Gold Antifade reagent containing DAPI (Thermo Fisher Scientific). Sections were digitized with a Zeiss Axiovision Scanner (Zeiss) and analyzed using HALO v3.2.1851.354 and the HighPlex FL v4.0 algorithm to segment and count immunopositive and immunonegative cellular subpopulations (Indica Labs). Representative images of multiplex immunofluorescence from HTAPP-102-SMP-11 were acquired on a Zeiss LSM980 confocal microscope. Images were further processed using spectral unmixing on Zen Blue (Zeiss) to separate channels containing cross-excitation artifacts.

**Methylation profiling**

Data from the National Cancer Institute’s Therapeutically Applicable Research to Generate Effective Treatments (TARGET; <https://ocg.cancer.gov/programs/target>) initiative, phs000467, were used as reference data to generate our methylation groupings. For methylation of samples from the HTAPP cohort, genomic DNA was extracted from formalin-fixed paraffin embedded (FFPE) tissue using the Maxwell DNA FFPE kit. FFPE-derived DNA was then hybridized to Illumina MethylationEPIC BeadChip (850k) arrays according to the manufacturer’s instructions.

**Data analysis of methylation data**

Preprocessing steps were performed using the *minfi* package. We excluded samples with log median intensity less than 10 in both the methylated (M) and unmethylated (U) channels. We then used the noob background subtraction method for dye-bias normalization^54^ and performed subset-quantile within array normalization (SWAN)^55^, a within-array normalization correction for the technical differences between the Type I and Type II array designs. In addition, we filtered probes within the X or Y chromosomes and probes containing polymorphic nucleotides (Infinium Methylation manifest column, "PROBE_SNPS") within 10-50 base pairs of and including the targeted CpG site.

For dimensionality reduction and visualization, we performed principal component analysis (PCA) in the initial steps using the top 5000 most variable probes. t-SNE dimensionality reduction was performed using the top 50 PCAs with a perplexity value of 22 and 5000 iterations (*Rtsne* package)^16^. Clustering was performed using consensus clustering (*ConsensusClusterPlus* package)^17^. We identified the most differentially methylated regions with *DMRcate*^18^.

**NetBID hidden driver analysis of single-nucleus RNA-seq data**

For snRNA-seq data, we adapted the network-based integrative NetBID algorithm^28^ to identify cell-type specific drivers from the snRNA-seq data. We first reverse-engineered cell-type specific transcription factor and signaling molecule interactomes using SJARACNe^29^, an information theory-based algorithm for regulatory network inference. Then, the *cal.Activity* function in NetBID was employed to infer the activities of each driver candidate in each nucleus from their gene expression profiles and the corresponding network. Since snRNA-seq data are extremely sparse and noisy, we use unweighted mean activity. The unweighted mean activity of a hub gene *i* in nucleus was defined by the following equation:

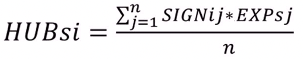

The gene expression matrix was z-normalized for each nucleus, where *EXPsj* is the expression value of gene *j* and *SIGNij* is the sign of Spearman correlation between gene *i* and its target gene *j*. The total number of targets for driver *i* is denoted by *n*. To identify drivers comparing group II MES or ADRN cells to the rest of the methylation groups, differential activity was calculated by using the *getDE.BID.2G* function in NetBID.

**CIBERSORTx analysis for bulk deconvolution**

We used CIBERSORTx^30^, a machine learning-based algorithm, to deconvolute bulk RNA-seq data of neuroblastoma using annotations from our snRNA-seq data. We down-sampled the snRNA-seq data to 10,000 nuclei to meet the computing requirement of CIBERSORTx. In the CIBERSORTx Cell Fractions Analysis Module, we used snRNA-seq reference matrix and bulk RNA-seq expression matrix of 131 neuroblastoma samples from the TARGET cohort (<https://ocg.cancer.gov/programs/target>) as input and used the default parameters. The proportions of each cell type identified from snRNA-seq data in each NBL bulk sample were reported.

**Multiplexed Ion Beam Imaging (MIBI) - Antibody Conjugation and Panel Assembly**

Lanthanide-labeled antibodies were conjugated using the Maxpar X8 Multimetal Labeling Kit (Fluidigm) according to the manufacturer’s protocol, with slight modifications^56^. For each antibody, 100 μg BSA-free antibody was washed and concentrated with conjugation buffer in an Amicon Ultra 50kDa filter (Millipore Sigma #UFC505096). Washed antibody was subsequently reduced with TCEP (Thermo Fisher Scientific #77720) at a final concentration of 4 μM for 30 min in a 37 ºC water bath. Reduced antibody was then mixed with lanthanide-loaded polymer with a functionalized maleimide group for 90 min in a 37 ºC water bath, followed by purification using an Amicon Ultra 50kDa filter. Conjugated antibody concentration was measured using a NanoDrop (Thermo Scientific) in IgG mode (280 nm). Final concentration was then adjusted to a suitable concentration (generally between 250-500ug/ml) with at least 30% v/v Candor Antibody Stabilizer (Thermo Fisher Scientific #NC0414486) before long-term storage at 4 ºC. Antibodies used in this study are listed in Table S8.

**MIBI Gold Slide Preparation and Treatment**

Gold slide preparation has been described previously^33,57–60^. Superfrost Plus glass slides (Thermo Fisher Scientific #12-550-15) were soaked in dish detergent diluted in double distilled water (ddH_2_O), then briefly wiped with Microfiber Cleaning Cloths (Care Touch #BD11945). Cleaned slides were then rinsed in ddH_2_O to remove any remaining detergent and air-dried with compressed air to remove any remaining moisture. Sputter deposition of 30 nm of tantalum and 100 nm of gold was performed by the Microfabrication Shop of the Stanford Nano Shared Facility.

To enhance tissue section adhesion, gold slides were treated with Vectabond Reagent (Vector Labs #SP-1800-7) according to the manufacturer’s instructions. Slides were first dipped in ddH_2_O, transferred into acetone for 5 min, before incubation with 1:50 diluted Vectabond Reagent in acetone for 10 min. Slides were then rinsed with ddH_2_O to quench and remove excessive reagents. Any remaining water was removed by gently wiping the slide with a Kimwipe, followed by air-drying.

**MIBI Staining Protocol**

The procedure for MIBI staining was previously described^33,57,59,61^. Gold slides with FFPE tissue sections were baked at 70 ºC for 1 hour, then incubated in xylene for 10 min twice, before a series of rehydration steps using an automated linear stainer (3 washes in xylene, 3 washes in 100% ethanol, 2 washes in 95% ethanol, 1 wash in 80% ethanol, 1 wash in 70% ethanol, 3 washes in ddH_2_O). Each wash was performed for 180 seconds. Slides, after completion of rehydration, were temporarily stored in ddH_2_O. Then, heat-induced epitope retrieval was performed using a pH 9 retrieval buffer (Dako #S236784-2) for 10 min at 97 ºC, followed by controlled cooling to 65 ºC, followed by further cooling of the slides in the retrieval tank on the bench for 30 min. A hydrophobic PAP pen was used to circle out the tissue sections to conserve reagents and antibodies. The slides were then blocked by BBDG (5% normal donkey serum, 0.05% sodium azide in 1X tris-buffered saline IHC wash buffer with Tween 20) for 1 hr in a humidity chamber at room temperature, before staining with the MIBI antibody cocktail at 4 ºC overnight. Subsequently, slides were washed and post-fixed with 2% glutaraldehyde and 4% paraformaldehyde in 1X PBS. After post-fixation, slides underwent a series of dehydration washes on the linear stainer (3 washes in 100 mM Tris pH 7.5, 3 washes in ddH_2_O, 1 wash in 70% ethanol, 1 wash in 80% ethanol, 2 washes in 95% ethanol, 3 washes in 100% ethanol. Each wash was performed for 30 seconds. The slides are then stored in a vacuum desiccator until acquisition.

**MIBI-TOF Imaging and Image Processing**

Datasets used in this study were acquired on a custom MIBI-TOF mass spectrometer equipped with an O_2_ duoplasmatron ion source (Alpha iteration). The running parameters are listed below:

- Pixel dwell time: 8 ms
- Image area: 400 μm x 400 μm
- Image size: 512 x 512 pixels
- Primary ion current: 3.5 nA on a built-in Faraday cup
- Number of depths: 1

Low-level processing of MIBI images, such as image extraction, background removal, and denoising, was performed using a customized version of the MIBI Analysis tools^33,62^. For tiled runs, a custom MATLAB script (MIBIStitch) and flat-field correction methods were used to stitch and correct individual FOVs into a final tiled image.

Cell segmentation was performed with a local implementation of deepcell-tf 0.6.0^34,35^. Histone H3 was used as the nucleus channel and a dummy (all-zero) image as the membrane channel. Signals from input channels were first capped at the 99.7^th^ percentile before input into the model. *model_mpp*=1.5 was used for optimized segmentation results.

Extracted features for cells were clustered based upon phenotypic markers for immune cells (CD11b, CD163, CD68, CD3, CD4, CD8, CD20 and CD45), tumor (CD56), and endothelium (CD31 and CD34) using FlowSOM and marker enrichment modeling (MEM)^36,37^ as previously reported^57,59^. Cellular neighborhoods were called using k = 20^63^. FOVs from one sample, HTAPP-130-SMP-91, were omitted from downstream neighborhood analysis because only tumor cells were identified in the 3 FOVs from that sample.

**Patch Level Analysis of NeuroBlastoma (PLANB)**

Tumor-stroma boundaries were manually annotated using CD56 and vimentin to identify tumor-rich and stroma-rich regions, respectively. A patch mask was generated from that annotation, and the microenvironment around each tumor was extracted stepwise towards the inside and outside of the defined boundary. Each ‘step’ area was defined as the 1 pixel-width extension/inclusion area generated by MATLAB functions: *imdilate()* and *imerode()* on the patches masks, for up to 75 pixels inwards and outwards of the boundary.

To avoid interference from neighboring tumor patches while also keeping the integrity of the microenvironment of surrounding tumor, expansion pixels that overlapped with other original tumor patches were omitted in each step. Subsequently, the marker expression level and cell phenotype fraction (number of pixels for each cell type as a fraction of the total number of pixels in each ‘step’ area) were extracted. For downstream analysis, the average expression level of each marker or cell type pixel percentage, along with the 95% confidence intervals, were plotted using the R package “ggplot”.

**Co-detection by indexing (CODEX) slide processing**

Multiplexed CODEX analysis of neuroblastoma tissues was performed using а panel of antibodies conjugated to custom DNA barcodes and detector oligos (see Table S8) as well as common buffers, robotic imaging setup and instructions for CODEX staining of frozen specimen from Akoya Biosciences. Other than pretreatment specific to FFPE tissues, these instructions and buffer compositions correspond to the procedure described previously in Schürch, et al.^63^ Frozen tissue was embedded in Tissue-Tek optimal cutting media (OCT) and frozen according to standard procedures. Blocks were sectioned on a cryotome into seven-micron sections after OCT blocks were equilibrated to the cryostat temperature for at least 30 minutes. Tissue sections were dragged over the surface of cold poly-L-lysine coated coverslips and spread inside the cryostat by transiently warming up the bottom surface of the coverslip with a finger before storage at -70 °C for 1-2 months.

**CODEX slide staining and data processing**

Section pre-processing was done as described previously in Golstev, et al.^39^ Prior to staining, sections removed from the freezer and dried for 5 min on a surface of Drierite. Dried coverslips were then dipped for 10 min into room temperature acetone, then fully dried for 10 min at RT. Sections were then rehydrated for 5 min in S1 buffer (5 mM EDTA, 0.5% w/v bovine serum albumin, and 0.02% w/v sodium azide in PBS), and further re-fixed for 20 min at room temperature in S1 buffer with 1.6% formaldehyde. Formaldehyde was washed off twice with S1 buffer, and sections were then equilibrated in S2 buffer (61 mM NaH_2_PO_4_∙7 H_2_O, 39 mM NaH_2_PO_4_ and 250 mM NaCl in a 1:0.7 v/v solution of S1 buffer and doubly-distilled water; final pH 6.8-7.0) for 10 minutes. Sections were then blocked in blocking buffer^63^ for 30 minutes.

Automated image acquisition and fluidics exchange were performed using an Akoya CODEX instrument driven by CODEX driver software (Akoya Biosciences) and Keyence BZ-X710 fluorescent microscope configured with 4 fluorescent channels (DAPI, FITC, Cy3, Cy5) and equipped with a CFI Plan Apo λ 20x/0.75 objective (Nikon). Hoechst nuclear stain (1:3000 final concentration) was imaged in each cycle at an exposure time of 1/175 sec. Biotinylated CD39 - detection reagent was produced as previously described^64^, used at a dilution of 1:500, and visualized in the last imaging cycle using DNA streptavidin- PE (1:2500 final concentration). DRAQ5 nuclear stain (1:500 final concentration) was added and visualized in the last imaging cycle. Each tissue was imaged with a 20x objective in a 7x9 tiled acquisition at 1386x1008 pixels per tile and 396 nm/pixel resolution and 13 z-planes per tile (axial resolution 1500 nm). Images were subjected to deconvolution to remove out-of-focus light.

Acquired images were pre-processed (alignment and deconvolution with Microvolution software (<http://www.microvolution.com/>) and segmented (including lateral bleed compensation) using publicly available CODEX image processing pipeline available at <https://github.com/nolanlab/CODEX>^39^.

**Slide-SeqV2**

Fresh frozen neuroblastoma tissue resections, embedded in optimal cutting temperature compound (OCT), were allowed to reach -20°C inside a cryostat (Leica, CM1950) for 30 min before handling. The tissues were mounted on a chuck with OCT and sliced at 10um thickness and a 5° cutting angle into tissue sections. Slide-SeqV2 pucks were generated as described previously in Stickels, et al.^40^ The Slide-SeqV2 puck was securely fastened to a microscope glass slide with a small drop of water with the beads facing upwards. The glass was turned upside down to ensure the puck was not moving during handling of the glass. With the puck facing down the puck surface was aimed at the region of interest in the tissue by lowering the puck over the tissue section and allowing a quick melting of tissue and puck to occur before removing the puck:tissue outside the cryostat. The puck was moved with forceps to a microcentrifuge tube pre-filled with 200 μl hybridization buffer (6x SSC with 2 U/μl Lucigen NxGen RNase inhibitor) and incubated for 15 min. The puck was then washed by dipping it once into 1x Maxima RT buffer.

The puck was moved to a second tube with 200 μl First strand synthesis mix (115 μl water, 40 μl Maxima 5x RT buffer, 20 μl of 10 mM dNTPs, 5 μl Lucigen NxGen RNase inhibitor, 10 μl of 50 μM template switch oligonucleotide (IDT, 5’ AAGCAGTGGTATCAACGCAGAGTGAATrG+GrG 3’) and 10 μl Maxima H Minus reverse transcriptase and incubated at room temperature for 30 min followed by 52°C for 90 min.

200 ul of Tissue digestion buffer (100 mM Tris-Cl pH 7.5, 200 mM NaCl, 2% SDS, 5 mM EDTA and 1:50 proteinase K) was added directly to the first strand solution, and the mixture was incubated at 37 °C for 30 min. The beads were thereafter removed from the cover glass surface by vigorously pipetting multiple times in the presence of 200 μl wash buffer (10 mM Tris pH 8.0, 1 mM EDTA and 0.01% Tween-20) and the glass removed by using forceps and discarded. The beads were pelleted by centrifugation at 3000g for 2 min and the supernatant removed. The bead pellet was resuspended in 200 μl of wash buffer and centrifuged again and this was repeated for a total of three washes.

After the last wash the beads were resuspended in 200 μl of Exonuclease I mix (170 μl water, 20 μl ExoI buffer and 10 μl Exonuclease I) and incubated at 37 °C for 50 min. 200 μl of wash buffer was added directly to the exonuclease mixture and centrifuged for 2 min at 3000g. Following supernatant removal from the bead pellet, the wash step was repeated for a total of three times. The bead pellet was resuspended in 200 μl of 0.1N NaOH and incubated for 5 min at room temperature. 200 μl of wash buffer was added and the beads were centrifuged for 2 min at 3,000g and the wash repeated for a total of three times.

Next, second-strand synthesis was performed on the beads by addition of 200 μl of second-strand synthesis mix (133 μl water, 40 μl Maxima 5x RT buffer, 20 μl of 10 mM dNTPs, 2 μl of 1 mM dN-SMRT oligonucleotide and 5 μl Klenow enzyme) to the bead pellet and incubation at 37 °C for 1 h. After second-strand synthesis, 200 μl of wash buffer was added directly to the mixture and centrifuged for 2 min at 3000g. Following supernatant removal from the bead pellet, the wash step was repeated for a total of three times. The bead pellet was then washed one further time with 200 μl water followed by resuspension of the bead pellet in 50 μl PCR mix (22 μl water, 25 μl of Terra Direct PCR mix buffer, 1 μl of Terra polymerase, 1 μl of 100 μM TruSeq PCR handle primer and 1 μl of 100 μM SMART PCR primer). PCR was performed according to the following program: 95°C for 3 min; four cycles of 98°C for 20 s, 65°C for 45 s and 72°C for 3 min; 13 or 14 cycles of 98°C for 20 s, 67°C for 20 s and 72°C for 3 min; 72°C for 5 min; hold at 4°C.

The PCR product was purified with 30 μl (0.6x volumes) of AMPure XP beads and incubated 10 min at room temperature. The magnetic beads were then pelleted using a magnetic separator for 5 min followed by two washes with 80% ethanol for 30 sec each and the cDNA eluted with 50 μl EB solution. The bead purification was then repeated once more at a 0.6x volume of beads:cDNA and final elution was done with 12 μl EB. The size and concentration of the final cDNA was assessed on a Bioanalyzer High Sensitivity DNA chip (Agilent) and on a Qubit high sensitivity dsDNA kit (Invitrogen).

600 pg of sample cDNA was converted into Illumina compatible sequencing libraries through tagmentation with a Nextera XT kit according to the manufacturer’s instructions. The tagmented libraries were indexed via PCR amplification with TruSeq5 (IDT, 5’- AATGATACGGCGACCACCGAGATCTACACTCTTTCCCTACACGACGCTCTTCCGATCT-3’) and N700 series barcoded index primers with the PCR program: 72 °C for 3 min; 95°C for 30 s; 12 cycles of 95°C for 10 s, 55°C for 30 s, 72°C for 30 s and 72°C for 5 min; hold at 4°C.

The sequencing ready libraries were purified with AMPure XP beads at a 0.6:1 volume ratio beads:DNA as described earlier and eluted with 12 μl EB solution. The size and concentration of the final sequencing libraries were assessed on a Bioanalyzer High Sensitivity DNA chip (Agilent, 5067-4626) and on a Qubit high sensitivity dsDNA kit (Invitrogen, Q32851), respectively. Finally, the library concentration was normalized to 4 nM for sequencing and pooled together. All 19 samples were sequenced on an Illumina NovaSeq S2 flow cell 100 cycle kit. The sequencing settings were: read 1- 42 bases, read 2 - 41 bases and index 1- 8 bases for the i7 index.

**Slide-SeqV2 data analysis**

We generated gene expression matrices for the Slide-seqV2 bead barcodes using the Slide-Seq tools alignment pipeline, with bcl files from the raw Slide-SeqV2 data as the input^40^. Within Seurat, for each tissue section, we filtered out cells with fewer than 50 UMIs detected and cells outside a radius of 2550 pixels^9^. We then carried out clustering analysis using the Seurat functions *SCTransform*, *RunPCA*, *FindNeighbors*, and *FindClusters* with default values. To build the *k-*NN graph, as input we used the first thirty principal components, and to perform Louvain clustering on the graph, we used a resolution parameter of 0.3. The clusters were primarily used to guide the assignment of tumor and stroma regions described below.

Because Slide-SeqV2 data does not have single-cell spatial resolution and beads can capture transcripts from multiple cells, cell type assignment at each bead requires deconvolution using a reference dataset of purified cell types. As a cell type reference, we used all cells from our annotated scRNA-Seq and snRNA-Seq data, collected from the same ten patients profiled by Slide-SeqV2. Due to the count sparsity of Slide-SeqV2 data, we used annotation at the level of broad cell lineages: ADRN, sympathoblast, MES, myeloid cell, T/NK cell, B cell, stroma, endothelial cell. To perform the deconvolution of each Slide-SeqV2 bead, we used the RCTD algorithm, which outputs the inferred proportion of each cell type at each bead^41^. We ran the RCTD algorithm with doublet_mode flag set to “full”, as based on H&E images, we expected beads may capture transcripts from more than two cells. When visualizing the Slide-SeqV2 beads mapped to their spatial coordinates or performing co-localization analysis, we assigned to each bead the cell type that had the greatest inferred proportion by RCTD. To identify pairs of cell types that are significantly colocalized and may be interacting, we used the squidpy package to build a spatial neighbors graph and quantify neighborhood enrichment^43^. A positive neighborhood enrichment score indicates a pair of cell types are more spatially proximal than expected by chance, and conversely, a negative neighborhood enrichment score indicates a pair of cell types are less spatially proximal than expected by chance.

To increase both the gene throughput and accuracy in our Slide-SeqV2 experiments, we used the Tangram method to integrate the Slide-SeqV2 data with scRNA-Seq data^42^. Tangram aligns the single cell profiles in space to maximize spatial correlation of the gene expression of the mapped single cells and of the spatial data. For each spatial sample, we mapped the single cell data collected from the same patient. As training genes, we first identified the top fifty marker genes (Mann-Whitney U test) for each of the 10 broad cell lineages (ADRN, sympathoblast, MES, myeloid cell, T/NK cell, B cell, stromal cell, endothelial cell, Zona glomerulosa, erythrocyte) across all the combined scRNA-Seq and snRNA-Seq data. We then filtered this gene set to only those genes detected in both the single cell and spatial datasets, and that had mean number of counts > 0.001. When mapping the single cells into space, we trained for 1000 epochs with a learning rate of 0.1 and an RNA count based prior. After completing the alignment, we used the mapped data to examine predicted spatial gene expression patterns.

We expected that a specific cell type’s gene expression would be different in the tumor vs stromal regions of the tissue. To assign spatial regions as tumor or stromal, within each cell cluster we summed the inferred cell type proportions across all beads from RCTD, with the simplifying assumption that there are roughly a similar number of cells sampled by each bead. If the summed cell type proportions in a cluster were >50% malignant (ADRN, sympathoblast, MES), we assigned the cluster as ‘tumor-rich’, and otherwise as ‘stroma-rich’. We then applied the statistical model CSIDE, which allows us to detect genes within a specific cell type that are differentially expressed across different spatial contexts, while accounting for the effect that each bead has different cell type compositions^44^. We set the “cell_type_threshold” of 100 and set the FDR control to 0.1.

**QUANTIFICATION AND STATISTICAL ANALYSIS**

**Statistical analysis**

Bayesian compositional modeling of cell populations in Figures 2, 4, 5, and S4 were performed using scCODA using default parameters.^4^ Composition analysis was performed by sequentially using each cell type as a reference, and then majority voting was used to define credibly changed cell types. For all scCODA analyses in this study, the false discovery rate (FDR) cutoff was set to 0.05.

To compare sc-/snRNA-seq dataset parameters in Figures 3F-G, S1C-E, and S5B, a Wilcoxon rank-sum test was performed using the base R statistics command ‘wilcox.test’. To identify cell-subset specific marker genes, we tested for differentially expressed genes between a given cell subset and all other cells using a two-sided Mann-Whitney U test with an

FDR control at 0.05.

**ADDITIONAL RESOURCES**

- Data portal for all raw and processed data within this study: <https://humantumoratlas.org/>
- Interactive visualizer of integrated single-cell/nucleus RNA-sequencing data:

<https://viz.stjude.cloud/community/human-tumor-atlas-network-consortium~6>

**DATA AND CODE AVAILABILITY**

All sequencing data (bulk RNA-sequencing, whole exome sequencing, single-cell/nucleus RNA-sequencing) will be made available upon publication through the Human Tumor Atlas Network (HTAN) Data Coordinating Center portal (<https://humantumoratlas.org/>). This portal houses both raw data (fastq or bam files), processed data (count matrices or variant annotation files), and sample metadata tables. In addition to sequencing data, the HTAN data portal contains all highly multiplexed spatial proteomic data (from MIBI and CODEX) as well as Slide-SeqV2 data (raw and processed data with spatial coordinates). Processed single-cell/nucleus RNA-sequencing data can be interrogated using an interactive visualizer (<https://viz.stjude.cloud/community/human-tumor-atlas-network-consortium~6>). Methylation data from Illumina MethylationEPIC bead arrays will be accessible through GEO (accession GSE216650) upon the time of publication.
